# A high-throughput UAV millimetric imaging pipeline for plant density phenotyping reveals high heritability in wheat-pea sole crops and intercrops

**DOI:** 10.64898/2026.10.09.757598

**Authors:** V. Freitas, J. Enjalbert, T. Flutre, J.-M. Gilliot

## Abstract

Within-field diversification, notably through cereal legume intercropping, is a major lever of the agroecological transition. Breeding cultivars adapted to intercropping requires major changes in breeding practices, owing to strong genotype-by-stand-type interactions and the high spatial heterogeneity of low-input field trials. Part of this heterogeneity stems from variation in plant density, a first yield component that strongly influences resource competition and is critical to monitor in intercrops. Given the many plots in such platforms, high-throughput phenotyping of plant density is essential. Over two seasons, we designed a large field trial involving 200 wheat cultivars observed as sole crop as well as intercropped with two pea tester cultivars, in a pseudo-randomized complete block design. Plant density was manually counted on 40 cultivars, alongwith UAV flights at both high and low altitudes. We propose a method to predict plant density from millimeter-resolution RGB images, using linear regression on log-transformed fractional vegetation cover and additional colorimetric and textural indices. We predict total density of wheat or pea in sole crops, or the combined pea+wheat total density in intercrops with relative RMSE of 19.2% and 27.4% depending on season. Thanks to the pipeline, total plant density was predicted across the entire trial, comprising more than 1,000 microplots (200 varieties); this density showed high heritability (0.75-0.85). We provide guidelines regarding the composition of the training set for such intercrop trials. High-throughput phenotyping approaches such as the one proposed here can be instrumental in mobilizing genetic diversity for agroecology.

**PLAIN LANGUAGE SUMMARY:** Growing two crops together in the same field such as wheat and pea is a promising way to make farming more sustainable. But breeding varieties suited to this practice remains difficult, because many interacting factors shape how the two crops perform together. One key factor is the number of plants that actually emerge in each plot: it reveals how many plants may ultimately contribute to yield, and it strongly affects how plants compete for resources. Counting plants by hand across thousands of small plots is impossible, so we used a drone to take very detailed photos of a large experiment involving 200 wheat varieties, grown alone or mixed with pea. From these images, we developed a method to estimate the number of plants in each plot wheat, pea, or both. Measuring this trait early and across hundreds of varieties can help breeders develop crops adapted to more diverse, environmentally friendly farming systems.

## 1. INTRODUCTION

Modern crop production faces converging pressures from climate variability, soil degradation, biodiversity loss, and the need to reduce synthetic inputs (Tilman et al., 2002). This has motivated agroecological strategies based on crop diversification to maintain productivity while improving resilience and environmental performance (Beillouin et al., 2021). Cereal-legume intercropping is a prominent example, providing well-documented benefits, notably through improved resource use efficiency for nitrogen which allows maintaining total yield while reducing nitrogen fertilizer requirements (Bedoussac et al., 2015; E. S. Jensen et al., 2020; Rodriguez et al., 2020). However, the prevalence of genotype-by-stand type interactions (Demie et al., 2022) calls for breeding varieties better adapted to intercropping (Bančič et al., 2024; Salomon et al., 2026; Sampoux et al., 2020). In this goal, typical field trials aim at screening the performance of numerous genotypes (dozens in Moutier et al. (2022), hundreds in Salomon et al. (2026)) by cultivating them both as sole crops and as binary intercrops, most often combining a single genotype per species in each mix but see Haug et al. (2023). To handle the combinatorial explosion of the number of possible intercrops differing in genotype composition and sowing densities, several strategies were proposed, ranging from incomplete, yet balanced designs involving similar numbers of genotypes of both species (Haug et al., 2021, 2023), to tester-based designs in which many genotypes of a focal species are intercropped with a few ‘tester’ genotypes of the companion species (Moutier et al., 2022). However, even with such designs, field trials can still consist of hundreds or even thousands of micro-plots.

To quantify the relative productivity of genotypes across crop types, varieties are typically evaluated in terms of resource use efficiency in intercrop versus sole crop using land equivalent ratio (LER). The partial LER of a given variety corresponds to the surface needed in sole crop to obtain the same yield as in intercrop, and is calculated as yield in intercrop divided by yield in sole crop. The sum of pLERs gives the total LER. Partial LER is well known to depend on relative plant density (Yu et al., 2016), as is total yield well-known to depend on total plant density (Postma et al., 2021; Weiner & Freckleton, 2010). Accurate characterization of plant density is therefore critical for both agronomy and breeding purposes, and allows performance corrections when spatial heterogeneity is present (S. M. Jensen et al., 2023). We hence focussed here on measuring *total* plant density as a first step, keeping the measurement of *relative* plant density (i.e., per species in intercrop) for future work.

Conventional approaches rely on manual plant counts in quadrats or along short row segments (Whaley et al., 2000). This method is labor-intensive and only provides a sparse spatial sampling incompatible with large field trials. Ground-based imaging operated by a human (e.g., handheld or tripod-mounted cameras) can alleviate part of this burden but remains limited in throughput and area coverage at the scale of field phenotyping campaigns (Araus & Cairns, 2014; Sankaran et al., 2015). Uncrewed aerial vehicles (UAVs) equipped with RGB or multispectral sensors, now provide an operational platform for high-throughput, field-based phenotyping (Gano et al., 2024; Jin et al., 2021; Sankaran et al., 2015). UAV imagery offers rapid coverage of large experimental areas, flexible acquisition timing and potentially very high spatial resolution. It has been successfully employed to estimate plant density and other structural traits in low-density crops such as maize and sunflower, where individual plants are clearly separable (Gnädinger & Schmidhalter, 2017; Olson et al., 2024). In contrast, estimating plant density in dense canopies such as wheat, where individual plants overlap even at early stages, remains challenging.

In the literature, there are two main categories of methods for counting plants using UAV imagery: (i) direct methods that involve the explicit extraction of individual plants using classical techniques of segmentation, mathematical morphology, and connected component analysis; (ii) indirect methods that estimate plant density based on vegetation cover properties, without attempting to extract individual plants. Varela et al. (2018) obtained an accurate count of the number of corn plants using a direct method. In a recent review dedicated to plant counting (Pathak et al., 2022), the authors showed that these direct approaches are primarily applicable to row crops such as corn, soybeans, cotton, or sugar beets. Object-based image analysis (OBIA) is an approach for detecting objects using both spectral and spatial information. (Nuijten et al., 2019) used a region-growing algorithm to create multi-pixel object primitives based on spatial and spectral features, on endive crops. When objects are extracted, geometric parameters are often calculated to help in plant identification. Koh et al. (2019) developed an algorithm based on template matching for safflower seedling detection, using 1000 image patches (templates) representing various plant examples.

When plants are close together and there is potential for leaf overlap, individual counting becomes difficult or even impossible, so indirect approaches are preferred. In the case of wheat at emergence, Jin et al. (2017) demonstrated that there was a strong relationship between the observed density and the vegetation cover fraction derived from UAV images acquired at very low altitude. Various variables derived from UAV images can be used to predict plant density. It can be done by measuring fractional vegetation cover (FCOVER) and related vegetation indices, as they have been shown to correlate well with plant density in dense canopies of cereals cultivated as sole crop (Jin et al., 2017; Wilke et al., 2021). Some authors succeeded in improving the accuracy of plant density predictions by incorporating variables that account for image texture. For example, Shen et al. (2024) used gray-level cooccurrence matrix (GLCM) texture features (Haralick et al., 1973) in their study of corn density. The use of texture information is less common in the literature for plant counting than for estimating aboveground biomass (Y. Liu et al., 2022; Yin et al., 2025). Various modeling approaches were used to predict plant counts. Regression-based methods that link FCOVER to density provide computationally efficient and easy to deploy alternatives. However, these previous works have been restricted to a few tens to approximately one hundred micro-plots (Jin et al., 2017; Wilke et al., 2021), and were not tested at the scale of a complete breeding platform which, in contrast, can include several thousand micro-plots. The transfer of these methods to large-scale high-throughput phenotyping hence represents a significant challenge. Moreover, with the introduction of breeding efforts for intercropping, it is also necessary to assess total plant density in platforms containing micro-plots sown with different species in sole crops and the corresponding intercrops. Handling such an heterogeneity remains largely unexplored. Machine learning-based models such as Partial Least Squares Regression (PLSR), Support Vector Machine (SVM), Random Forest, and XGBoost are currently used in phenotyping applications, as for measurement of photosynthetic traits (Lu et al., 2025) but are rarely or never used for plant counting. Deep learning-based models has shown promise for counting individual plants or organs in dense canopies (Aich et al., 2018; Peng et al., 2022; Yang et al., 2025) but practical deployment is constrained by the need for large annotated training sets, as well as substantial computational cost during training (Osco et al., 2021; Wang et al., 2025).

Here, we develop and validate an approach to predict total plant density in both dense and sparse canopies of cereal-legume intercrops and their corresponding sole crops using UAV-derived from FCOVER and other vegetation indices. Our strategy was to combine i) a high-altitude flight to survey the entire field trial and generate a georeferenced orthomosaic, using a photogrammetric approach, and ii) a low-altitude flight to acquire one high-resolution image per micro-plot to better discriminate soil vs. plant pixels, followed by the georeferencing of each low-altitude image across the orthomosaic. We then (i) computed FCOVER at high spatial resolution using a rule-based pipeline, in parallel, textural were derived directly from the raw input images, we also introduced object-based variables which were computed from the mask generated by FCOVER pipeline, the goal here is not to extract an entire plant object, but rather homogeneous image regions often referred to as “blobs”, and then calculate geometric properties used in predictions; (ii) calibrated and validated regression models relating FCOVER and other predictors to total plant density across a range of cereal-legume intercrops and sole crops differing in terms of sowing densities; (iii) assessed whether a single, generic model can be applied across stand types or whether stand-specific calibration is required; and (v) assessed the importance of genetic variations for total plant density at the scale of the whole breeding platform.

## 2. MATERIALS AND METHODS

### 2.1 Plant material

The plant material consisted of 200 winter wheat varieties (focal species) and 2 winter pea varieties (considered as testers), thereafter referred to as the “MBD panel”. A subset of 40 wheat varieties and one pea variety will be referred to as the “CB panel”. The wheat varieties were classified into five earliness groups, with group 5 being the earliest and group 1 the latest. Each group had the same size: 40 genotypes per group for MBD, and 8 genotypes per group for CB.

### 2.2 Field trial

A field experiment was conducted during the cropping season of 2023-2024 (season 1) and replicated in 2024-2025 (season 2), at the “Le Moulon” site of the Versailles-Saclay experimental unit (UEVS) on the Saclay plateau (Gif sur Yvette, France; 48.7155, 2.1512), on two different experimental fields (Figure 1).

**Figure 1:**
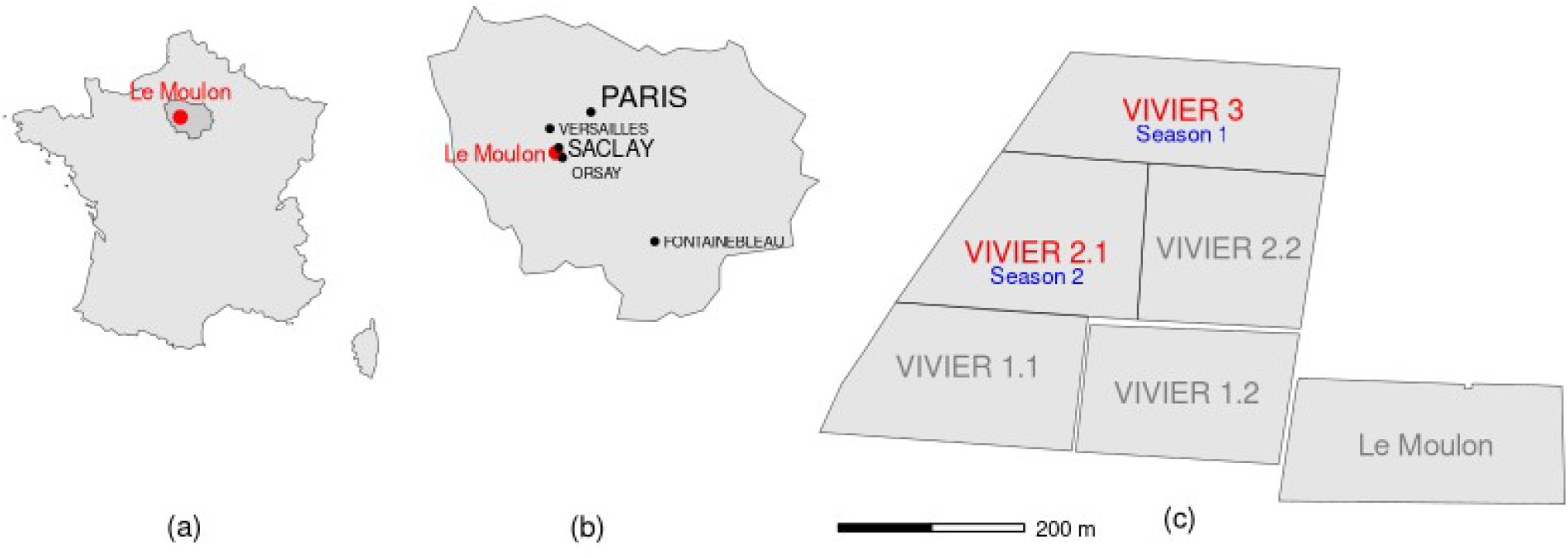
location of the field trials, (a): France, (b) Paris region (Ile de France), and (c) “Le Moulon” site of INRAE Versailles-Saclay experimental unit.

Both field trials followed a pseudo-randomized complete block design with two blocks. Each block was subdivided into 7-meter long x 1.50-meter wide micro-plots, sowed with 7 rows separated by 17.7 cm. The width used in the analyses below hence was 1.24 m, corresponding to the 1.06-m-wide seed drill in addition to a half-inter-row on both sides (Figure 2). Each micro-plot was sown either as sole crop of wheat at 300 seeds.m^-2^ (SCW), sole crop of pea at 40 seeds.m^-2^ (SCP), or intercrop of wheat at 150 seeds.m^-2^ and pea at 40 seeds.m^-2^ (ICWP).

**Figure 2:**
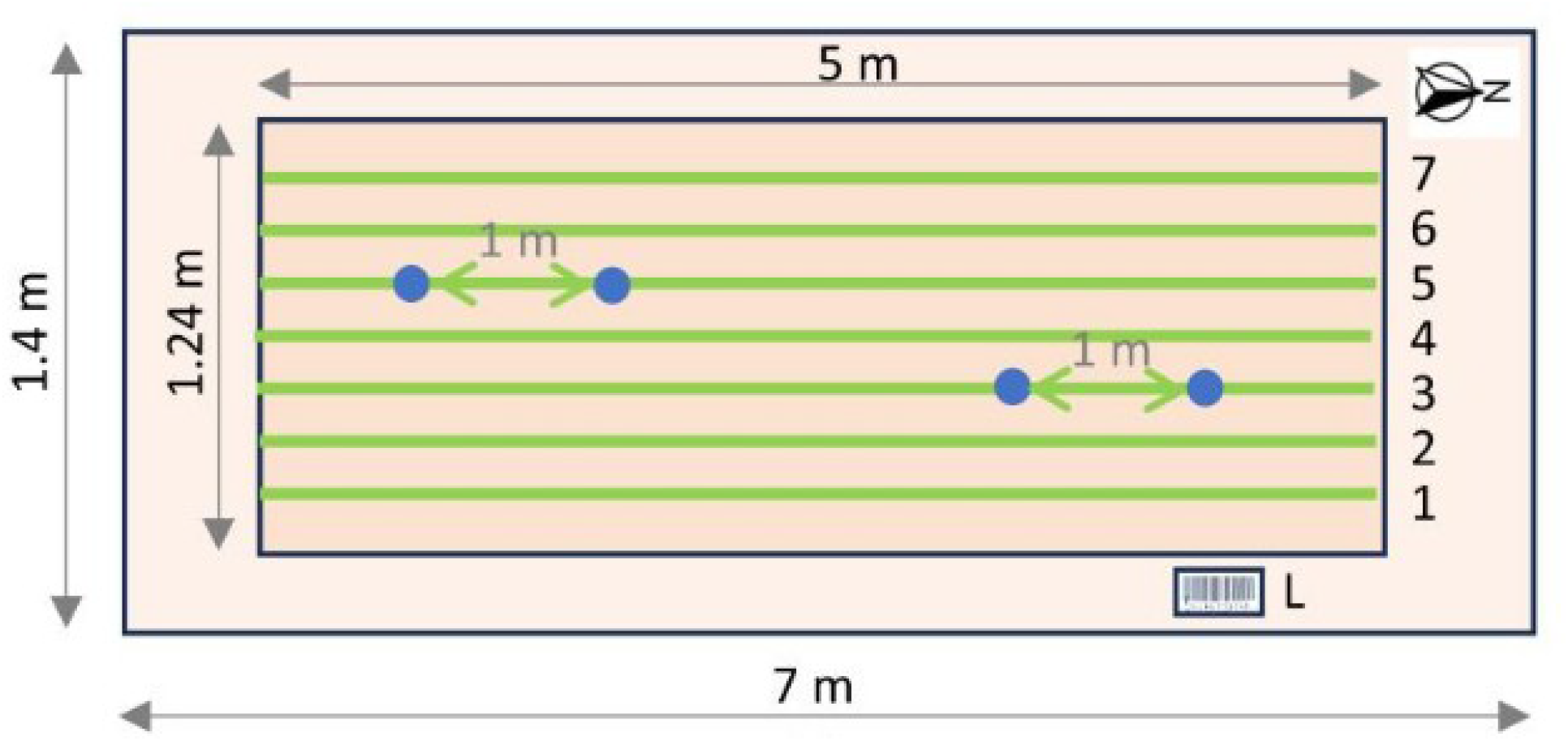
scheme of a micro-plot with 7 sowing rows, and both linear meters inside (between two blue dots), along which plants were manually counted. L was a barcode sticker used to identify the micro-plot in the field.

The field trial in season 1 comprised a total of 1795 micro-plots: 1310 labeled as MBD and used for grain yield assessment, and 170 labeled as CB and used for biomass sampling (the rest being extra and border micro-plots, Figure 3A). Similarly, the field trial of season 2 comprised a total of 1893 micro-plots: 1310 for MBD and 190 for CB (the rest being extra and border micro-plots, Figure 3B). The micro-plots were arranged in a regular grid, numbered from east to west (“X” field) and from north to south (“Y” field), a field named “XY” was created and used as an identifier of the micro-plots, by concatenating their X and Y coordinates, with for example, “100_12” corresponding to X=100 and Y=12.

**Figure 3:**
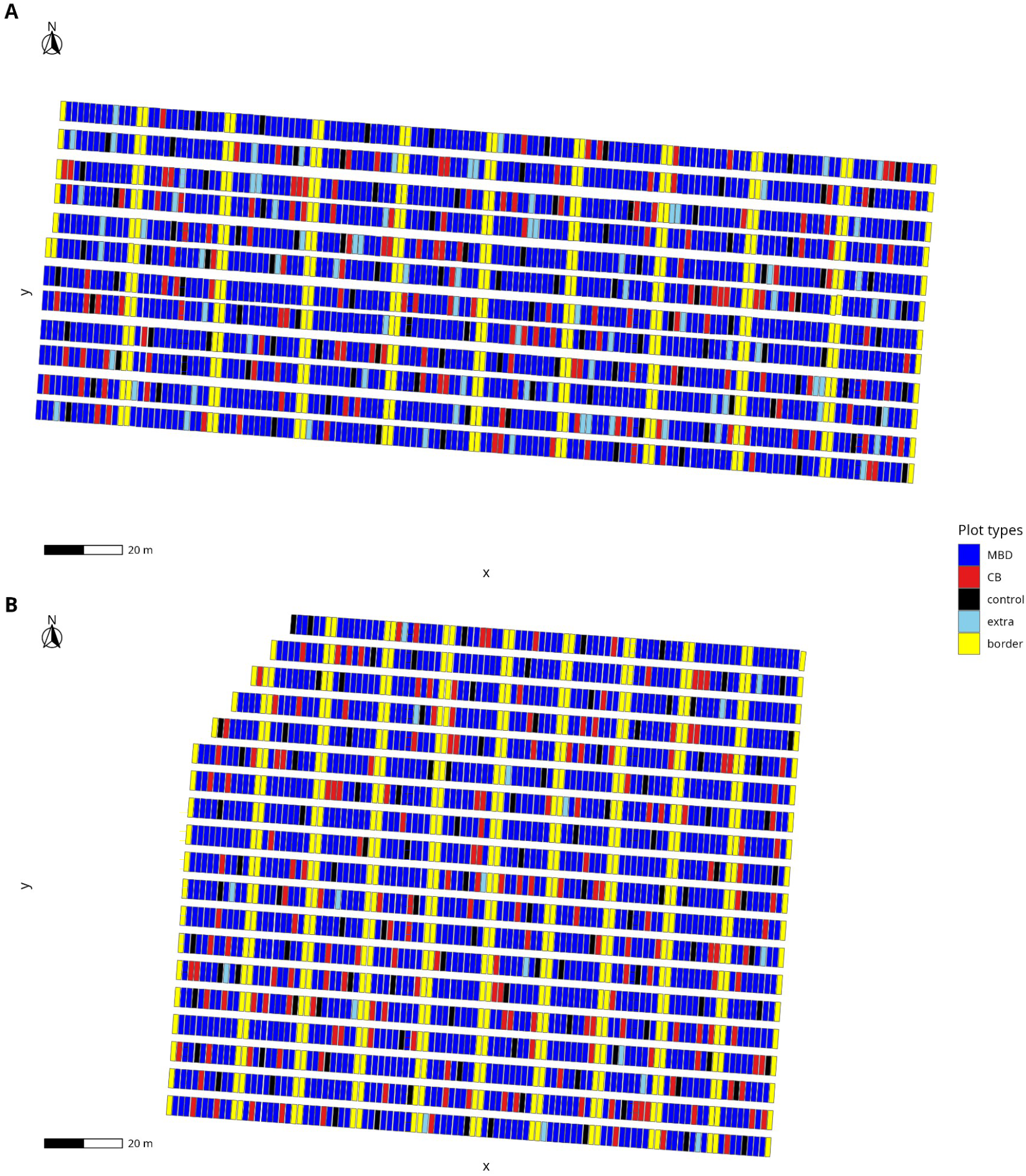
spatial layout of the field trials in season 1 (A) and season 2 (B). Colors indicate the micro-plot types, “CB” corresponding to the micro-plots in which plant density was manually scored along linear meters (two per micro-plot).

Sowing was done on November 24 and 25, 2023 for season 1 and November 4 and 5, 2024 for season 2. A single pesticide treatment took place the day after sowing (pre-emergence herbicide applied to limit weed development). Post-winter nitrogen residues reached 40 kg N.ha ^-1^ on season 1 (respectively, 27 kg N.ha^-1^ on season 2), a mineral fertilizer application was carried out in May at a rate of 40 kg N.ha^-1^ in season 1 (resp., in April at a rate of 30 kg N.ha^-1^ in season 2).

### 2.3 Data collection

#### 2.3.1 In field manual seedling counts

Ground-truth, manual seedling counts were performed on CB micro-plots only. Among the 170 CB micro-plots of season 1, 80 were in sole crop (40 per block), 80 were intercrops (also 40 per block) and the remaining 10 were controls (five per block). The same design was used in season 2, with 30 controls instead of 10 (15 per block). Each season (on 02/05/2024 for season 1 and on 12/16/2024 for season 2), two linear meters were delimited in each of the 160 non-control micro-plots, one on row 3 towards the northern side, and the other on row 5 towards the southern side (Figure 2), resulting in a total of 320 linear meters (160 in SC and 160 in IC). Each linear meter was delimited by a cyan-colored stake on each side (Supplemental Fig. S1). Plant counts (PC) were manually scored, in the field, per linear meter, then converted into plant densities (PD, plants.m^-2^) using equation 1:

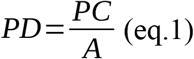

where A was the surface of the sampled linear meter (1 m long x 0.177 m wide).

#### 2.3.2 UAV-based image acquisition and processing

##### Sensors

Aerial imagery was acquired using a DJI Phantom 4 RTK (P4-RTK; DJI, Shenzhen, China), a lightweight quadcopter (1.39 kg) equipped with a centimeter-level GNSS real-time kinematic (RTK) positioning system. It carried an integrated 20-megapixel RGB camera (5472 × 3648 pixels) with a 13.2 × 8.8 mm CMOS sensor, 8.8 mm focal length, and an 84° field of view (Figure 4).

**Figure 4:**
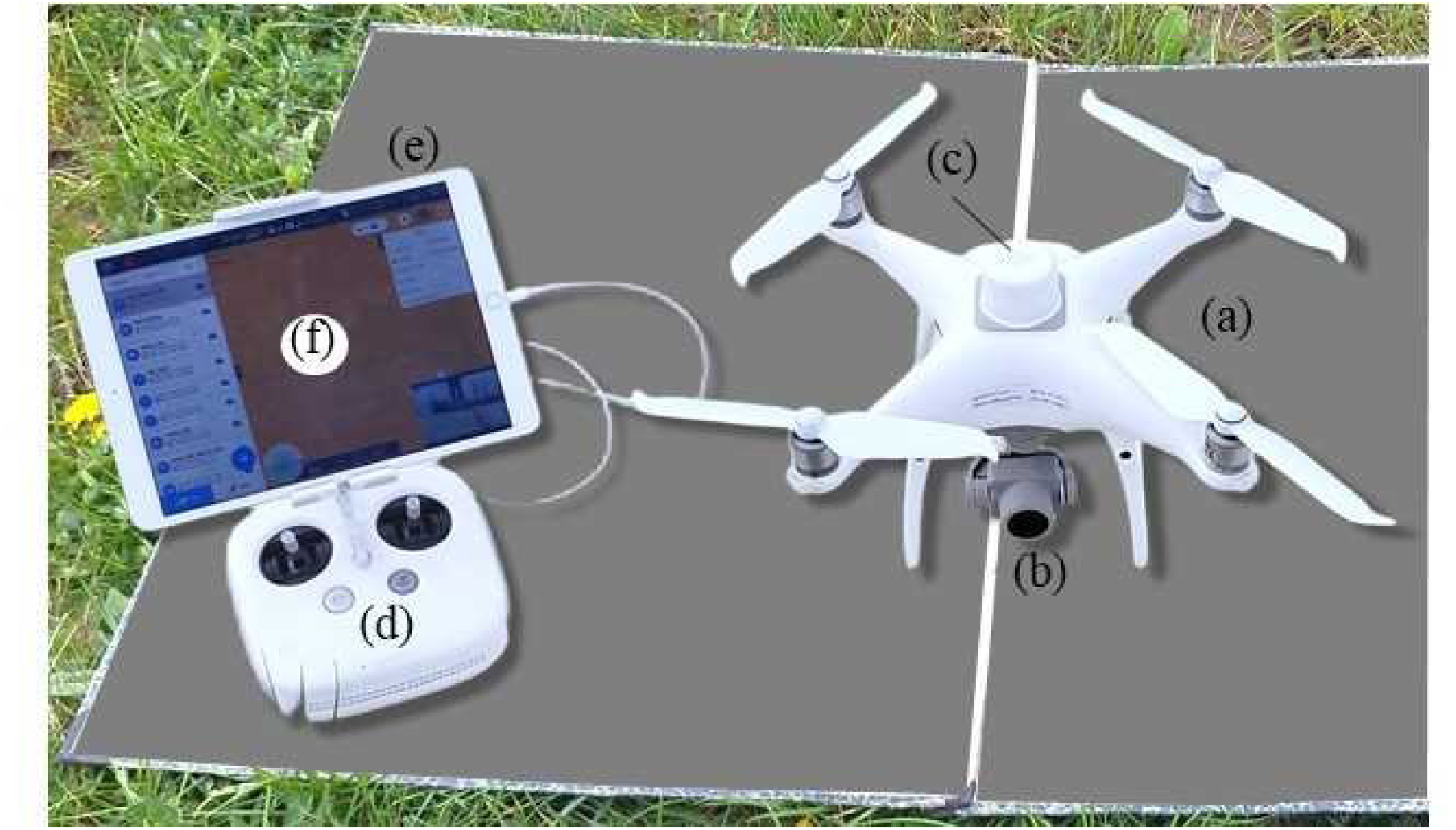
DJI Phantom 4 RTK with (a) UAV, (b) RGB camera, (c) RTK antenna, (d) GS SDK controller connected to (e) iPad tablet using (f) GS PRO application.

##### Flight configurations

Flights were performed at two different altitudes (Table1): (i) high-altitude flight at 25 m above ground level (AGL) with a ground sample distance (GSD) of 0.69 cm, and (ii) low-altitude flight (5 m or 6 m AGL) with a GSD of 0.16 cm. Flight missions were planned using Ground Station Pro software (GS PRO from DJI). Waypoint Route mode (Hover & Capture) was used for low altitude flights: to avoid blurry images, the UAV paused for one second on each plot. KML files containing the exact plot center positions were imported from QGIS software, to ensure precise alignment with the experimental design. For high-altitude flights, GS PRO was used in 3D map area mode, and an image overlap of 80% forward and 80% side was applied. All images were acquired from a nadir view and were scheduled, whenever possible, to avoid strong wind (< 15 km.h⁻¹) and variable cloud cover. Nonetheless, some flights were carried out under higher wind speed or intermittent cloud cover. The ISO was set to 100, and images were stored in JPEG format. Four flights were analyzed in this work, at 6 meters on 02/14/2024 and 25 meters on 02/28/2024 for season 1, and at both 5 meters and 25 meters on 12/20/2024 for season 2, all micro-plots being phenotyped at both altitudes.

### 2.4 Image processing and feature extraction

#### 2.4.1 Orthomosaic reconstruction from 25m flights

Images from the 25m flights were processed using a photogrammetric approach with Agisoft Metashape software (Agisoft LLC, St. Petersburg, Russia) to produce a georeferenced orthomosaic of the entire experimental site. The workflow included: (1) photo alignment, (2) dense point cloud generation, (3) digital elevation model (DEM) construction, and (4) orthomosaic reconstruction. The coordinate reference system used was Lambert 93. Real time kinematics (RTK) positioning ensured direct georeferencing without additional ground control points.

#### 2.4.2 Image georeferencing from low-altitude flights

Low-altitude flight images (5 or 6 meters high) were georeferenced at a first step, with a “direct” method, using: (i) the RTK GNSS positions, (ii) the three-axis attitudes (roll, pitch, yaw), and (iii) the UAV AGL elevation (derived from the 25m DEM). To further improve alignment, at a second step the direct images were georeferenced across the 25m orthomosaic. For season 1, a manual method was used, in QGIS (version 3.40), using the “Georeferencer tool”: ground control points (GCP) were selected in each micro-plot image and matched with corresponding positions on the orthomosaic. A first-order polynomial transformation (affine) was applied, and nearest-neighbor resampling was used to preserve the original pixel values. For season 2, a new procedure was performed, named “multi-scale image registration” (MUSIR; Gilliot et al, in prep) to automatically extract the GCPs. The Musir method relies on extracting characteristic points (keypoints) from images using Lowe’s algorithm (1999). The points are then matched between the high-resolution image and the orthomosaic, and their positions can then be used as GCPs. A first-order polynomial transformation (affine) was applied using these GCPs.

#### 2.4.4 Buffer definitions

Image analyses were performed on georeferenced rectangular areas, named “buffer” hereafter. A first set of buffers captured each linear meter, 1-m long and 16-cm wide (slightly less than the inter-row distance to avoid the inclusion of leaves from both neighboring rows). A second set of buffers matched each micro-plot, of 5-m long and 1.24-m wide.

#### 2.4.5 Canopy cover estimation

The fraction of vegetation cover (FCOVER) was estimated from the georeferenced images at both altitudes, using a multi-step raster processing pipeline implemented in R via the qgisprocess package (version 0.4.2; (Dunnington et al., 2024) which provides an R interface to the QGIS geoprocessing algorithms (version 3.44). First, red, green and blue channels (R, G and B; digital numbers between 0 and 255) were normalised into r, g and b (equation 2):

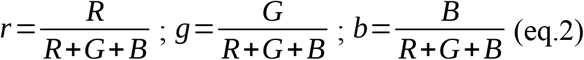

Second, the Excess Green vegetation index (equation 3) was computed pixel-wise for each buffer (the ones for the linear meters or for the micro-plots):

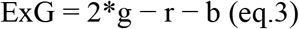

Third, the image was converted to the hue-intensity-saturation (HIS) colour space. The Intensity channel was used to trim under-exposed areas (shadow; but not over-exposed ones because of confounding with pea leaves). The Hue channel was used to detect cyan stakes used for manual seedling counts. Fourth, to discriminate vegetation from soil, Otsu’s method (Banerjee et al., 2021; Otsu, 1979) was tested and discarded as it overestimated vegetation cover due to a dominant soil fraction at this early growth stage (see (Jin et al., 2017). We thus developed a multi-criteria, thresholding method, applied in the exact same way to all micro-plots, so that a binary vegetation mask was generated by retaining pixels satisfying equation 4:

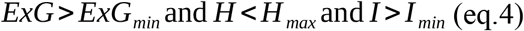

where *ExG_min_* = 0.08 was the ExG threshold (visually calibrated on a subset of sole crop wheat (SCW) micro-plots), *H_max_* = 100 (in QGIS-GRASS units, [0–255]) was used to exclude cyan-blue pixels (stakes), and *I _min_* = 60 (for season 1, 70 for season 2) was a lower bound on intensity used to discard shadowed pixels that would otherwise be erroneously classified as vegetation. For micro-plots of sole crop pea (SCP), a higher ExG threshold (*ExG_min_* = 0.11) was applied to account for the more reflective pea canopy that inflated FCOVER at the standard threshold (no change was needed for wheat-pea intercrops ICWP). Fifth, to remove residual artefacts (isolated noise pixels, small soil patches misclassified as vegetation), the binary mask was post-processed with the QGIS r.clump function, which finds regions of contiguous identical values and assigns a unique code to each pixel of the same region(labeling). These regions are then analyzed using r.stats.zonal function (count method) to determine their area, only regions with an area of at least 25 pixels (∼ 0.64 cm² at our resolution) were retained. This filtered binary vegetation mask preserved true plants while removing noise. Sixth, zonal statistics were then calculated within each buffer: the total valid pixel count (N_total_) and the vegetation pixel count (N_veg_), the latter being the sum of the previous binary mask. FCOVER was then calculated as in equation 5:

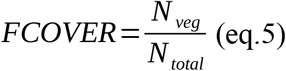

yielding one value per buffer. FCOVER was computed using these six steps for all buffers from all flights of both seasons.

#### 2.4.6 Morphological, spectral and textural descriptors

Beyond FCOVER, nineteen descriptors were extracted from each buffer to assess their potential of improving plant density prediction by capturing additional features from UAV images (Table 2). To ensure methodological consistency, all descriptors were derived from the same filtered binary vegetation mask used to compute FCOVER.

**Table 1:** configuration of UAV flights analyzed in this work (AGL: above ground level; GSD: ground sample distance).

| Date | Season | Height<br>AGL in m | GSD in<br>cm | Number of<br>flights | Flight<br>duration<br>(in hours) | Number<br>of images |
| --- | --- | --- | --- | --- | --- | --- |
| 02/14/2024 | season 1 | 6 | 0.16 | 5 | 1.98 | 1376 |
| 02/28/2024 | season 1 | 25 | 0.69 | 2 | 0.5 | 524 |
| 12/20/2024 | season 2 | 5 | 0.14 | 7 | 3.54 | 1536 |
| 12/20/2024 | season 2 | 25 | 0.69 | 3 | 0.94 | 478 |

**Table 2:** 19 predictors of plant density tested in this study.

| Group of predictors | Variable | Description |
| --- | --- | --- |
| 3 canopy cover predictors | FCOVER | Fraction of pixels with ExG |
|  | FCOVER_sd;<br>FCOVER_cv | standard deviation and coefficient of variation of FCOVER across buffers within micro-plot (used only for linear-meter buffers) |
| 4 blob detection predictors<br>(segmentation morphometry) | n_blobs_perM | Number of individually-segmented green blobs per square meter |
|  | mean_blob_area | Mean area of detected blobs |
|  | sd_blob_area | Standard deviation of blob area |
| | mean_compactness | Mean blob compactness ($4\pi \cdot A / P^2$) |
| 12 spectral descriptors | *Frac_003 ;...; frac_011* | Fraction of pixels with ExG > s, for $s \in \{0.03, 0.05, 0.07, 0.09, 0.11\}$ |
|  | *sd_r ; sd_g ; sd_b* | Standard deviation of red, green and blue channels within the buffer |
|  | *sd_VARI* | Standard deviation of VARI within the buffer |
|  | mean_ExG, sd_ExG, CV_ExG | Mean, standard deviation, and coefficient of variation of ExG within buffer |
| 60 texture descriptors<br>calculated on the<br>green channel | *glcm_contrast* | GLCM contrast |
|  | *glcm_entropy* | GLCM entropy (local heterogeneity) |
|  | *glcm_homogeneity* | GLCM homogeneity (texture uniformity) |
| (GLCM) 3 statistics<br>on 20 neo-channels<br>(5 window sizes: 5,<br>7, 9, 11, 15 pixels ×<br>4 directions: 0°,<br>45°, 90°, 135°) |  |  |

Morphological descriptors were obtained (EBImage R Package version 4.52, (Pau et al., 2010)) by connected component labelling of this mask within each buffer geometry, using the same algorithm at linear meter and plot scales. Shape descriptors were calculated using computeFeatures.shape function, which extracted objects (blobs) from the label image. Four descriptors were used: (i) the number of detected blobs (n_blobs), (ii) the mean blobs area (mean_blob_area), (iii) its standard deviation (sd_blob_area), and (iv) the mean compactness (mean_compactness = 4π·A / P², where A is area and P perimeter).

Spectral descriptors were computed from the ExG index and RGB rasters. Five ExG fractional indices (from frac_003 to frac_011) quantified the proportion of pixels exceeding increasing ExG thresholds (0.03, 0.05, 0.07, 0.09, 0.11) and provided a gradation of green biomass density independent of plant individuation. The standard deviations of normalized RGB channels (sd_r, sd_g, sd_b) and of the visible atmospherically resistant index (sd_VARI) were used to capture within-buffer spectral heterogeneity, with:

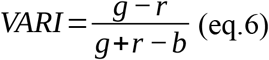

Other descriptors were the mean, standard deviation and coefficient of variation of ExG (mean_exG, sd_ExG, cv_ExG).

To test whether canopy texture could relate to plant density, texture features were derived from the grey-level co-occurrence matrix (GLCM; R package glcm version 1.6.6) on the green band, testing 20 configurations (five window sizes: 5, 7, 9, 11, 15 pixels, and four directions: 0°, 45°, 90°, 135°), each yielding three statistics (contrast, entropy, homogeneity), thus producing a total of 60 statistics.

### 2.5 Sequential selection of plant density predictors

The relation between FCOVER and manually-scored plant density was visually examined using locally weighted regression (loess, supplemental Fig. S4). Outliers were removed due to the presence of weed, waterlogging or sowing mistakes (13 cases in season 1 and 8 cases in season 2), leaving a total of 157 micro-plots in season 1 and 182 micro-plots in season 2 for model training and evaluation. The relationship between manually-scored plant density (PD) and the FCOVER as well as the descriptors listed above was then investigated in a two-step sequential manner using cross-validation. Negatively predicted plant densities were set to zero (such cases occurred for low densities in some sole crop pea micro-plots).

- In the first step, manually-scored plant density was linearly regressed on FCOVER only, using the base R function lm. Five transformations of FCOVER (log, square root, quadratic, cubic) were also evaluated individually, by R-repeated spatial K-fold cross-validation (with R=5 and K=10). “Spatial” refers to the fact that folds were stratified at the micro-plot level to ensure that the North and South buffers of the same micro-plot were always assigned to the same fold, hence preventing pseudo-replication. The performance metric used for comparison between the predicted values *X_i_* and the observed ones *O_i_* was the relative root mean square error (RRMSE, in %):

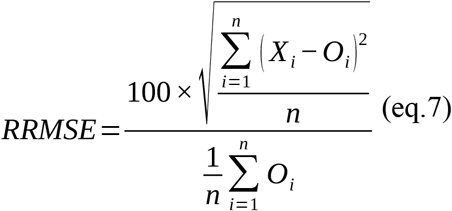

- In the second step, in addition to the transformed FCOVER, stand type (sole crop vs. intercrop) was introduced in the linear model. Two models were then compared: the first only having transformed FCOVER and stand type, and the second having an interaction term. A partial F-test (α = 0.05) determined which model was retained (with interactions for both years) using the base R function anova. The CV-RRMSE was obtained using the same R-repeated, spatial K-fold cross-validation as above.
- In the third step, 76 image-derived features were integrated using a regularized linear regression with the elastic net penalty for feature selection (R package glmnet version 4.1.10; (Friedman et al., 2010)), in addition to transformed FCOVER, stand type and their interactions that remained unpenalized (i.e., they were always selected). The overall prediction accuracy was assessed by R-repeated spatial K-fold cross-validation (with R=5 and K=10 as above) using the RRMSE. To optimize both components of the elastic net penalty (hyperparameters tuning), an inner k-fold cross-validation was performed (with k=10) using a squared-error loss function which is the default in glmnet for a Gaussian model. The mixing parameter *α* was optimized over the following sequence, seq(0, 1, by=0.1)^3^. For each *α* value, the penalty parameter *λ* was tested over a default sequence chosen by glmnet, and selected using the one-standard-error rule.

At the end of these three steps, the structure of the final model (the set of selected predictors, i.e., with non-zero coefficients) was obtained for each season, along with the corresponding CV-RRMSE. Other performance metrics were also computed: root mean square error (RMSE), bias and the coefficient of determination (R^2^). They were reported as mean ± standard deviation across the five repetitions.

### 2.6 Prediction accuracy across training sets with varying SC-vs-IC proportions

The influence of training on either sole crops only, intercrops only or both sole crops and intercrops, has been tested by varying the composition of the training set in a cross-validation scheme, and quantifying plant density prediction accuracy per target stand type (SC vs IC) with the same metrics as above (RRMSE, etc). A K=4 folds cross-validation was performed, repeated R=5 times. The size of the whole data set per season and stand type was n=160 linear meters, each fold hence being always of size 40, which left n_t_=120 for training and n_v_=40 for validation. Several training set compositions were compared: SC only (each fold was predicted after training on n_t_ sole-crop linear meters only, i.e 100% SC), IC only (n_t_ intercrop linear meters only) and xSC-yIC (n_t_ linear meters, x% of which being sole crops and the remaining y% being intercrops). Three SC-IC proportions were tested: 75% (75SC-25IC), 50% and 25%. More details are given in supplementary table S7.

### 2.7 Plant density prediction at the micro-plot scale for the whole breeding platform

For each season separately, the final model structures (one per season, c.f. section 2.5) were retrained on all linear-meter data from CB micro-plots (training set being SCW, SCP and ICWP) to estimate their parameters. Both models were then used to predict plant densities at the whole-micro-plot scale for all micro-plots (MBD + CB) of the breeding platform. The predicted densities on the CB micro-plots were compared to the manually-scored densities on the linear meters (averaged per micro-plot) through a linear regression for each season.

### 2.8 Wheat genetic variance estimation for plant density

The wheat genetic variance for plant density was estimated per season using data on all wheat genotypes (from both stand types, i.e., SCW and ICWP, to improve the size of the data set, hence estimation accuracy) using the following linear mixed model (LMM):

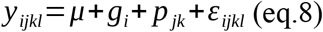

where y_ijkl_ is the manually-scored plant density of wheat genotype i (i in {1,…,I=40}) in block j (j in {1,J=2}), micro-plot k (k in {1,K_j_=2}) and linear meter l (l in {1,L_jk_}), μ is the intercept, g_i_ is the effect of genotype i modeled as random with *g_i_∼ N* (0 *, σ_w_*^2^) where σ_w_^2^ is the wheat genetic variance that is the focus of this analysis, p_jk_ is the effect of plot k in block j modeled as random to account for correlation between linear meters within the same micro-plot with *p_jk_ ∼ N* (0 *, σ_p_*^2^) where σ*_p_*^2^ is the micro-plot variance, and ε_ijkl_ is the residual error with *ε_ijkl_ ∼ N* (0 *, σ_e_*^2^). The manually-scored wheat plant densities in ICWP micro-plots were doubled beforehand to take into account the difference in wheat sowing density between stand types. This LMM was fitted using restricted maximum likelihood (R package lme4 version 1.1.37; (Bates et al., 2015). Broad-sense heritability was estimated using the following formula:

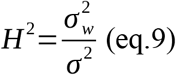

Because broad-sense heritability was computed on a genotype-mean basis (Falconer and Mackay, 1996), its denominator σ^2^ was estimated as:

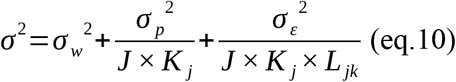

This model was then applied to manually-scored *total* plant densities (on CB micro-plots), with a correction applied to values from ICWP micro-plots: y_ijkl_^IC,corr^ = 2 c y_ijkl_^IC^. The factor “2” accounts for the difference in wheat sowing density in ICWP (150 seeds per m^2^) compared to SCW (300 seeds per m^2^). The correction factor “c” accounts for the fact that wheat plants only represent a subset of all plants in ICWP micro-plots. Instead of using the expected proportion of wheat plants, i.e.,150/190=0.79, we used the observed wheat proportion from manually-scored plant densities averaged over ICWP micro-plots: c=0.86 in season 1 and c=0.81 in season 2.

This model was finally applied to *predicted* total plant densities (on MBD micro-plots), with the same correction factor. The same formula applies for H2 except that L_jk_=1 in this case.

## 3. RESULTS

### 3.1 Data overview

#### 3.1.1 Manual plant count

Manual plant counts per species on a total of 340 one-meter linear buffers in 02/2024 for the season 1 (respectively, 380 in 12/2024 for season 2) were retained for analysis after quality control, corresponding to 170 micro-plots (resp., 190), distributed across three crop types (sole crop wheat, sole crop pea, and wheat-pea intercrop, supplemental Fig. S2). Both seasons were highly contrasted: the manual counts were centered on the expected counts in the second season but largely below for the first season (Figure 5).

**Figure 5:**
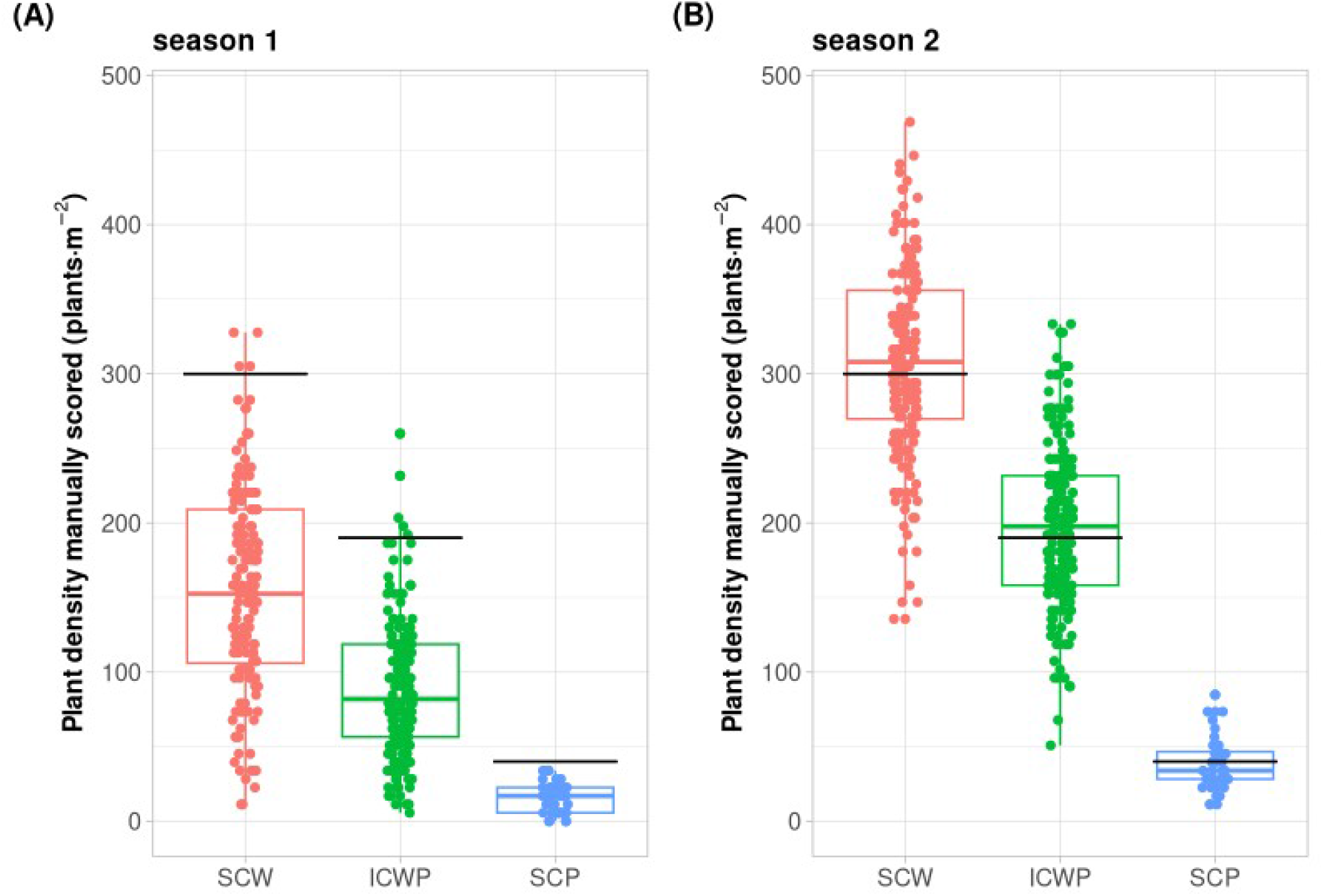
plant densities of each crop type (sole crop wheat SCW, sole crop pea SCP and wheat-pea intercrop ICWP) from manual counts in February 2024 for season 1 (A) and December 2024 for season 2 (B). Solid lines represent seed densities at sowing (in November for both seasons).

#### 3.1.2 Fcover computed from UAV images at 5-6m vs 25 m

UAV images differed between high-vs low-altitude in terms of resolution, with 0.69 cm at 25m compared to 0.14-0.16cm at 5-6m (Figure 6). FCOVER from 25m images was strongly underestimated in season 2, due to the lower contrast and images blurriness (Figure 6). As a result, the correlation between FCOVER at both altitudes hence was poorer due to this lower resolution at 25m (supplemental Fig. S3). We hence focused on the 5-6m images from now on.

**Figure 6:**
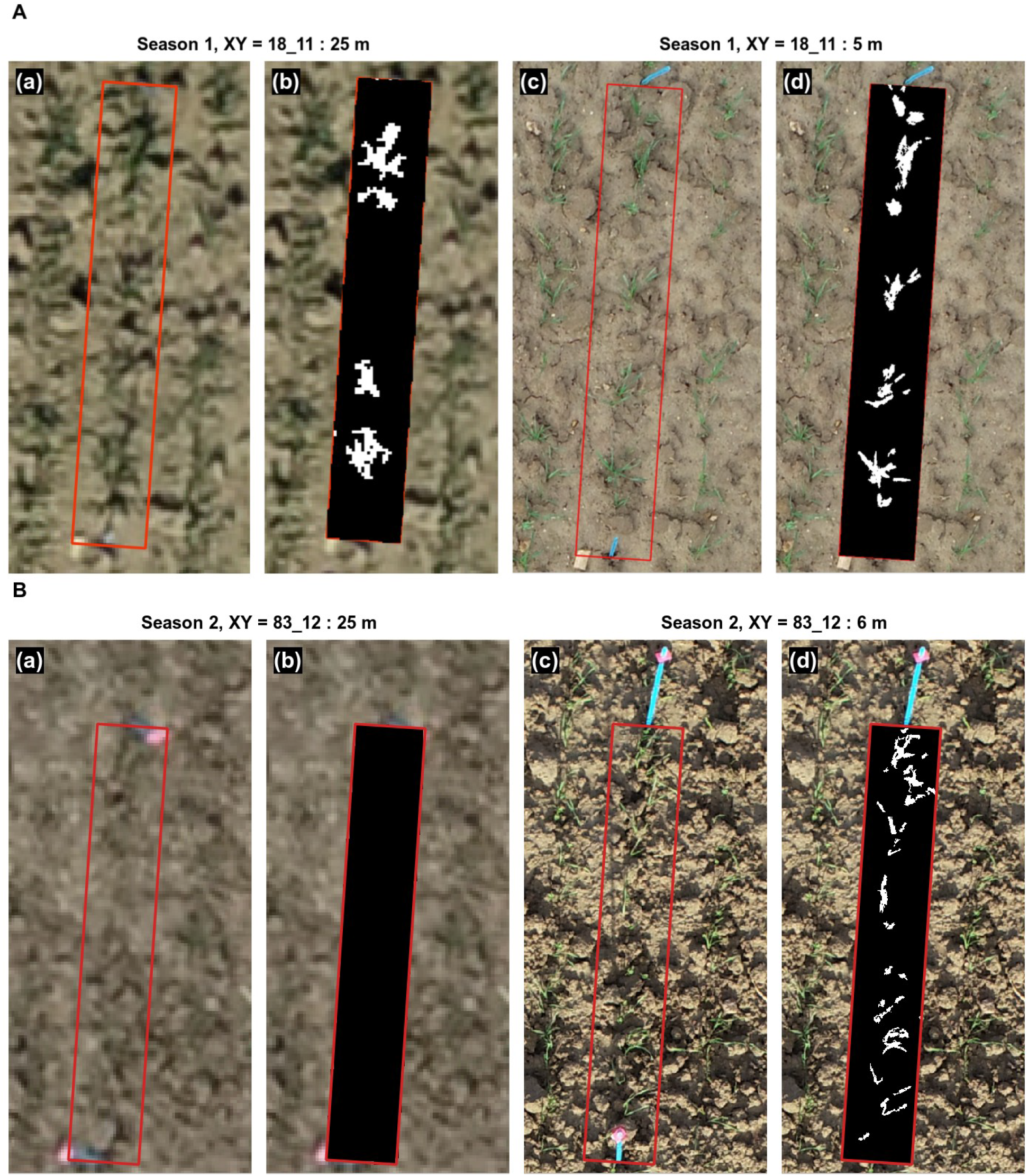
Example of UAV images centered on one linear-meter buffer (red rectangle), and results of Excess Green segmentation process (ExG) on this buffer, for both seasons and both flight altitudes: A) season 1, B) season 2, a) 6m RGB image, (b) ExG-based result on 6m image, (c) 25m RGB image, (d) ExG-based result on 25m image.

**Figure 7:**
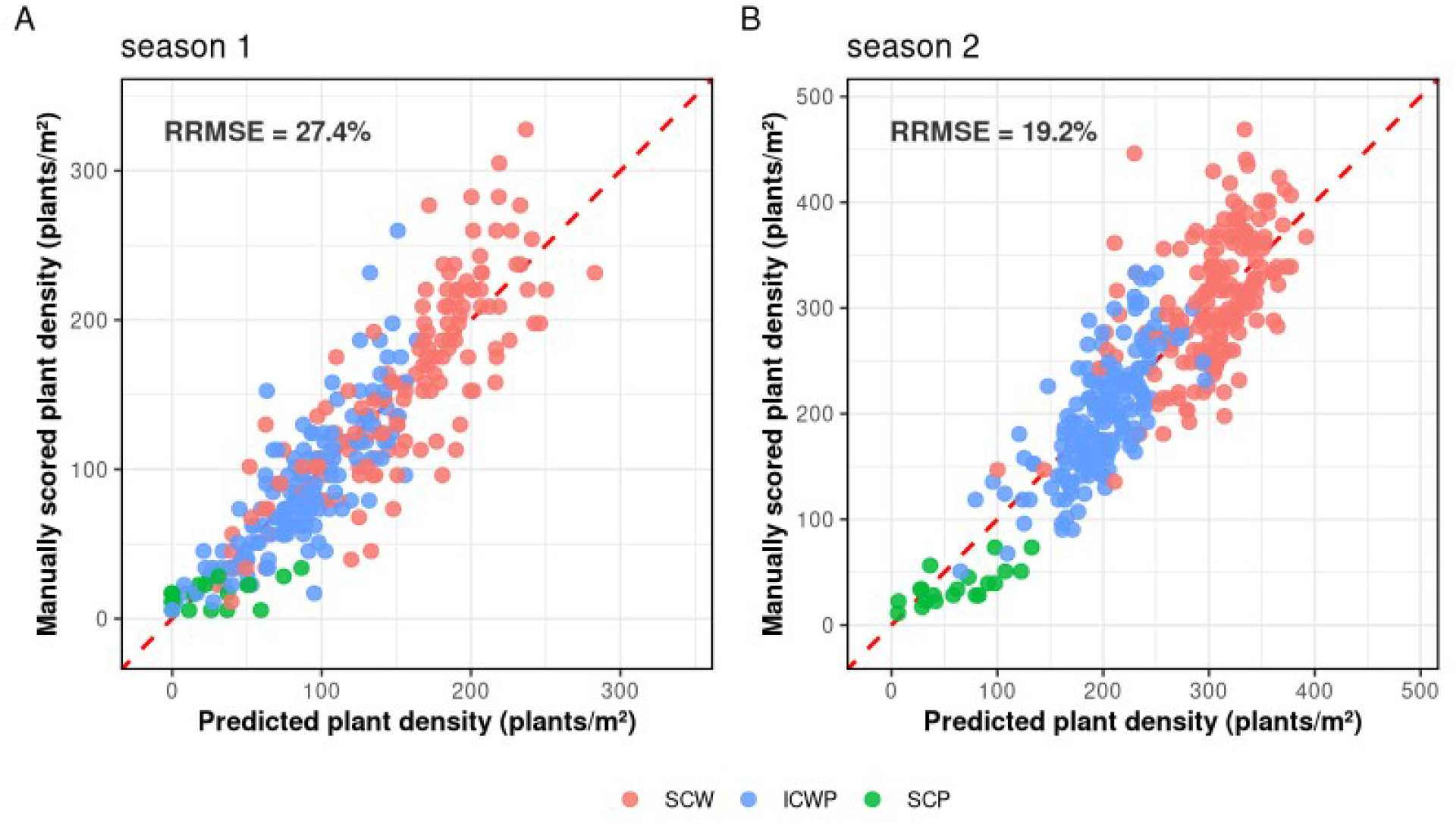
Comparison between manually-scored plant density and predicted plant density (averaged over five 10-fold crossvalidation replicates) from the final, season-specific models, for season 1 (A) and season 2 (B). Colors indicate the crop type (sole crop wheat SCW, sole crop pea SCP and wheat-pea intercrop ICWP).

In 5-6m images, median FCOVER was comparable between seasons (0.024 in season 1 and 0.056 in season 2). Moreover, it differed markedly between crop types. For instance in season 2, median FCOVER for sole crop wheat had the highest values (0.077), followed by wheat-pea intercrop (0.045) and sole crop pea (0.0045), consistent with the contrasting sowing densities and canopy architectures between species and crop types.

### 3.2 Prediction accuracy of plant density from RGB images

#### 3.2.1 Model selection via a sequential strategy

FCOVER was positively and strongly correlated with manual plant density across all linear-meter buffers (Pearson correlation coefficient: r = 0.699, p < 0.001, n = 314 in season 1; r = 0.680, p < 0.001 in season 2), confirming its relevance as a primary predictor.

Visual inspection of the FCOVER vs manual plant density relation revealed a non-linear pattern (Fig. S4). Several FCOVER transformations were hence compared by cross-validation based on the relative root mean square error (RRMSE): identity (i.e., no transformation), square root, logarithmic, quadratic and cubic. The logarithmic transformation outperformed the others in both seasons with a CV-RRMSE of 35.9% in season 1 (resp., CV-RRMSE = 25.4% in season 2, supplementary table S1). No combination of transformations improved upon the logarithmic alone (results not shown). The Pearson correlation coefficient between manual plant density and log(FCOVER) reached 0.80 in season 1 and 0.75 in season 2.

We then investigated the necessity to also account for differences in stand type (sole crop vs intercrop). Including in the model a “stand type” factor and its interactions with log-transformed FCOVER, significantly improved model performance for both seasons, with a CV-RRMSE of 31.5% in season 1 and 22.1% in season 2 (supplementary table S2).

Other spectral and texture features were included in a regularized model in addition to the transformed FCOVER, “stand type” factor and their interaction. The texture statistics were less consistently selected between the two years, with glcm_contrast more influential in season 1, while glcm_entropy was prevalent in season 2. Note however that 4 or the 5 glcm_contrasts (with an angle of 45°) were selected in both seasons. After hyperparameter tuning (supplemental table S3 and S5), the final model for season 1 reached a CV-RRMSE of 27.4% using 27 predictors, and 19.2% using 18 predictors for season 2 (supplemental table S4 and S6).

#### 3.2.2 Influence of training set composition

To quantify the influence of the training set composition, we trained on either sole crop-only data, intercrop-only data, or a mix of both with different proportions, and assessed prediction accuracy separately on sole crops and intercrops. Predictive performance depended on training composition (Figure 8). The lowest CV-RRMSEs were obtained when the test stand matched the training stand: 25.3% and 16.9% in sole crop or 36.3% and 19.6% in intercrops, for season 1 and 2 reciprocally. These CV-RRMSEs increased upon transfer to a different stand type, especially when predicting intercrop stands from sole-crop trained models (56.9% and 35.5% for season 1 and 2). In contrast, any training set composition with both SC and IC data (in proportions 75/25, 50/50 or 25/75) achieved similar precision across both targets in both seasons. CV-RRMSE remained between 22.1% and 38.1% in season 1 and between 15.6% and 21.6% in season 2, with absolute biases below 14 plants.m² in all cases. Differences between mixed compositions fell within the standard deviation of repeated cross-validations, indicating that the exact SC/IC ratio was not critical.

**Figure 8:**
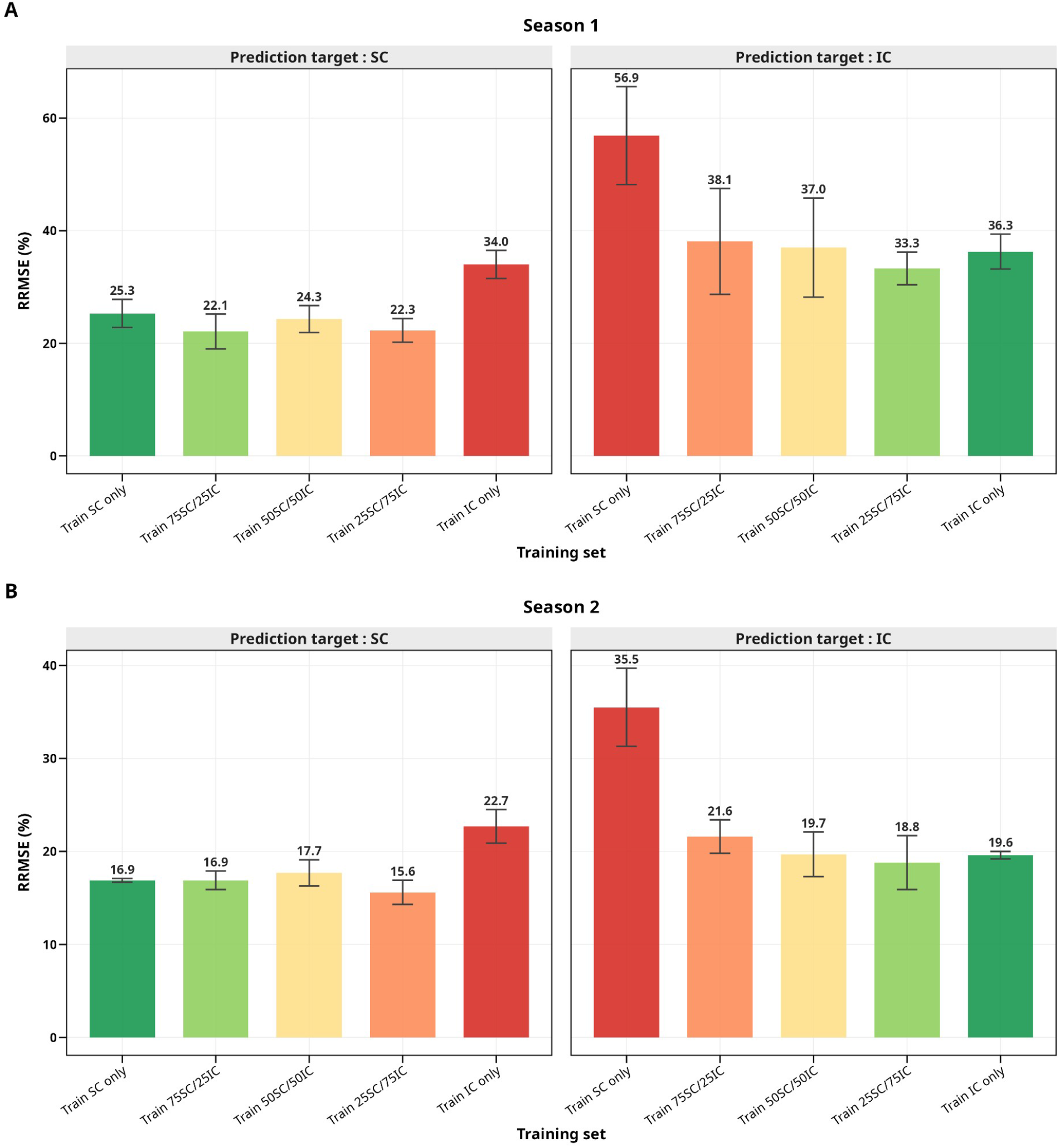
comparison of performance metrics in (A) season 1 and (B) season 2 for models trained on four different data sets contrasted for stand types. Colors indicate the relationship between training and prediction sets (green when both are of the same crop type, red when both are of different stand type, and along an orange gradient when training sets include both stand types).

### 3.3 Characterizing plant density at the scale of a whole breeding platform

The best, final model of each season, fitted on linear-meter data (see section 3.2.1), was used to predict plant density at the scale of the whole micro-plot (∼6.5 m²) for each of the ∼1200 micro-plots of the breeding platform.

To assess the relevance of these predictions, we first analyzed them for the CB micro-plots. Their comparison with the manual counting performed on their linear meter showed a strong agreement (RRMSE = 26.1% in season 1 and RRMSE = 17.7% in season 2; Figure 9).

**Figure 9:**
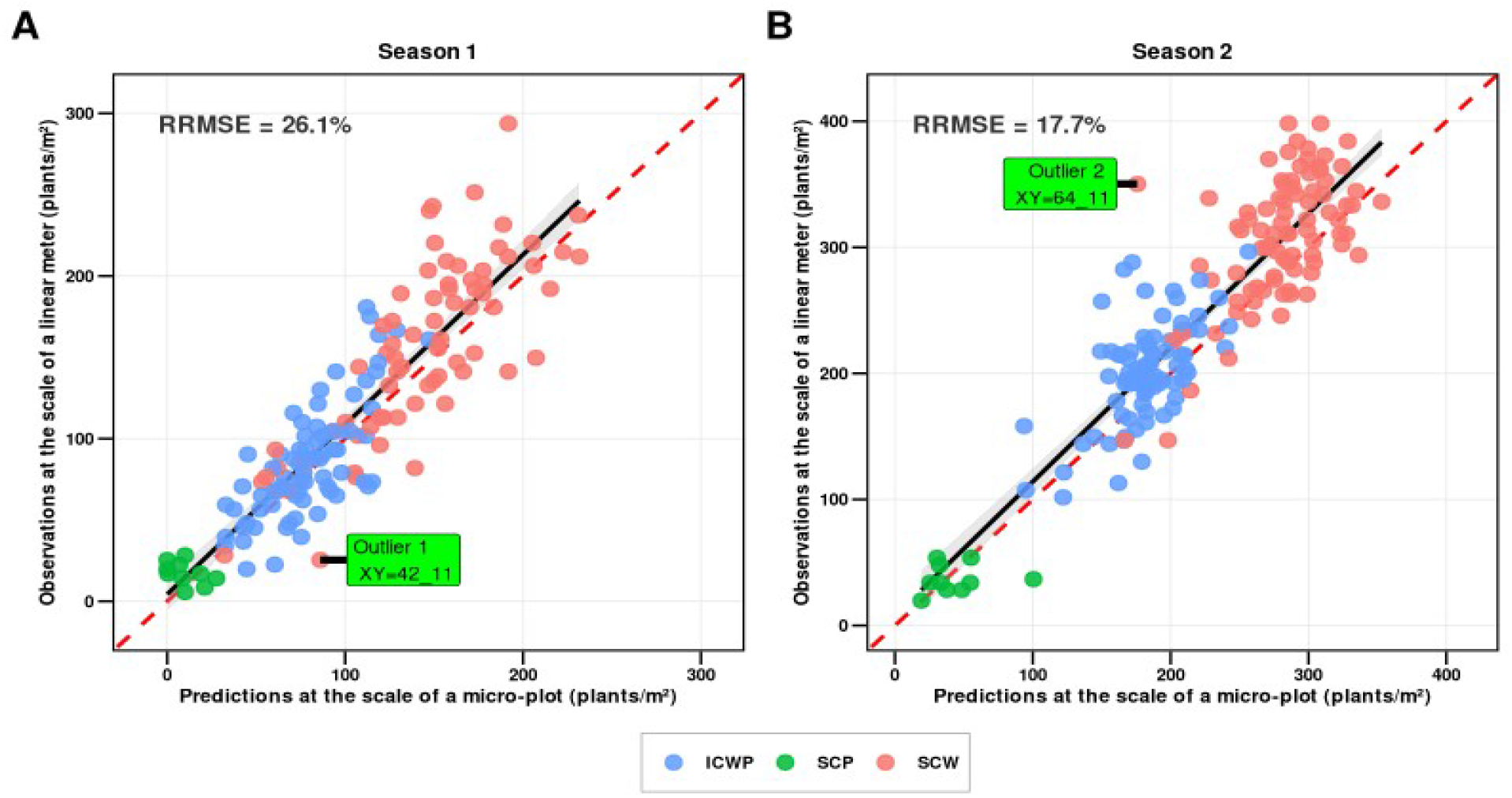
comparison of plant density predicted at the scale of a whole micro-plot versus plant density manually scored at the scale of a linear meter, for the CB micro-plots (observed linear-meter data were averaged per micro-plot). Colors indicate the crop type (sole crop wheat SCW, sole crop pea SCP and wheat-pea intercrop ICWP). Two outliers are highlighted (one per season).

The analysis of outliers in these correlations highlighted two main cases. Anomalous low plant densities in manually-scored linear meters (e.g., due to lower seed density or seed germination) led to underestimating plant density at the whole-micro-plot scale. Oppositely, the presence of holes in the micro-plot, outside of both linear meters, led to overestimating the plant density at the whole-micro-plot scale (Figure 10). These results highlight the interest of using whole-micro-plot images to have reliable plant densities despite the occurrence of within-micro-plot plant density heterogeneities.

**Figure 10:**
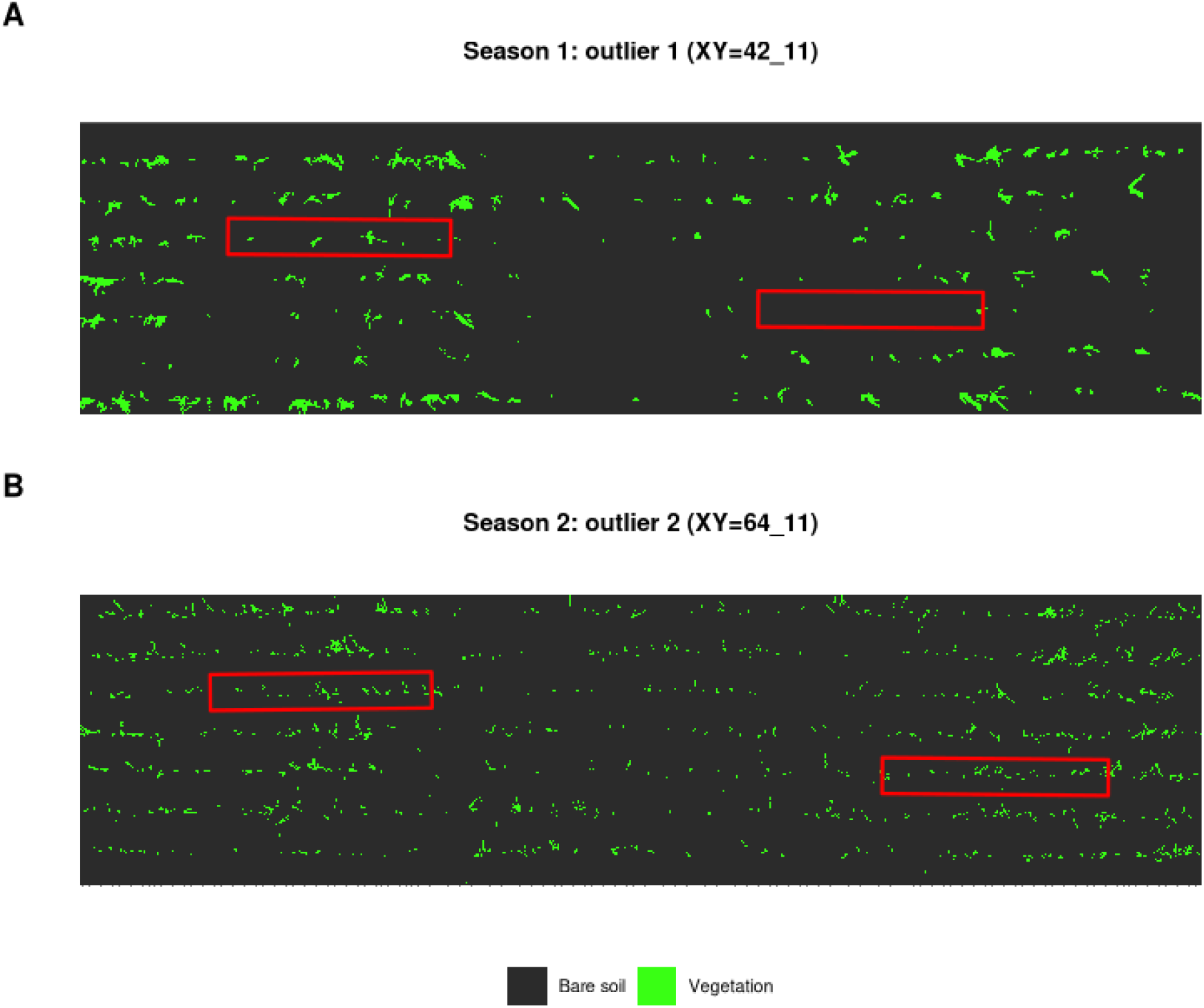
Analysis of 2 extreme outliers in the comparison of predicted plant densities of whole micro-plot versus manually-scored linear-meter plant densities (see Figure 9). A) Case of underestimation of micro-plot plant density (predicted at 86 plants.m-2) when using linear meter counts (averaged at 25 plants.m-2) in season 1. B) Case of over estimation in season 2 (predicted: 176, observed: 350). Scored linear meters are delimited by a red rectangle.

Using the fitted linear models, plant densities were predicted at the micro-plot scale on the whole MBD platform for each season. The distribution of predicted plant densities in both IC and SC were close to the ones estimated from manual counts (data not shown). The spatial distribution of these densities can be seen on Figure 11.

**Figure 11:**
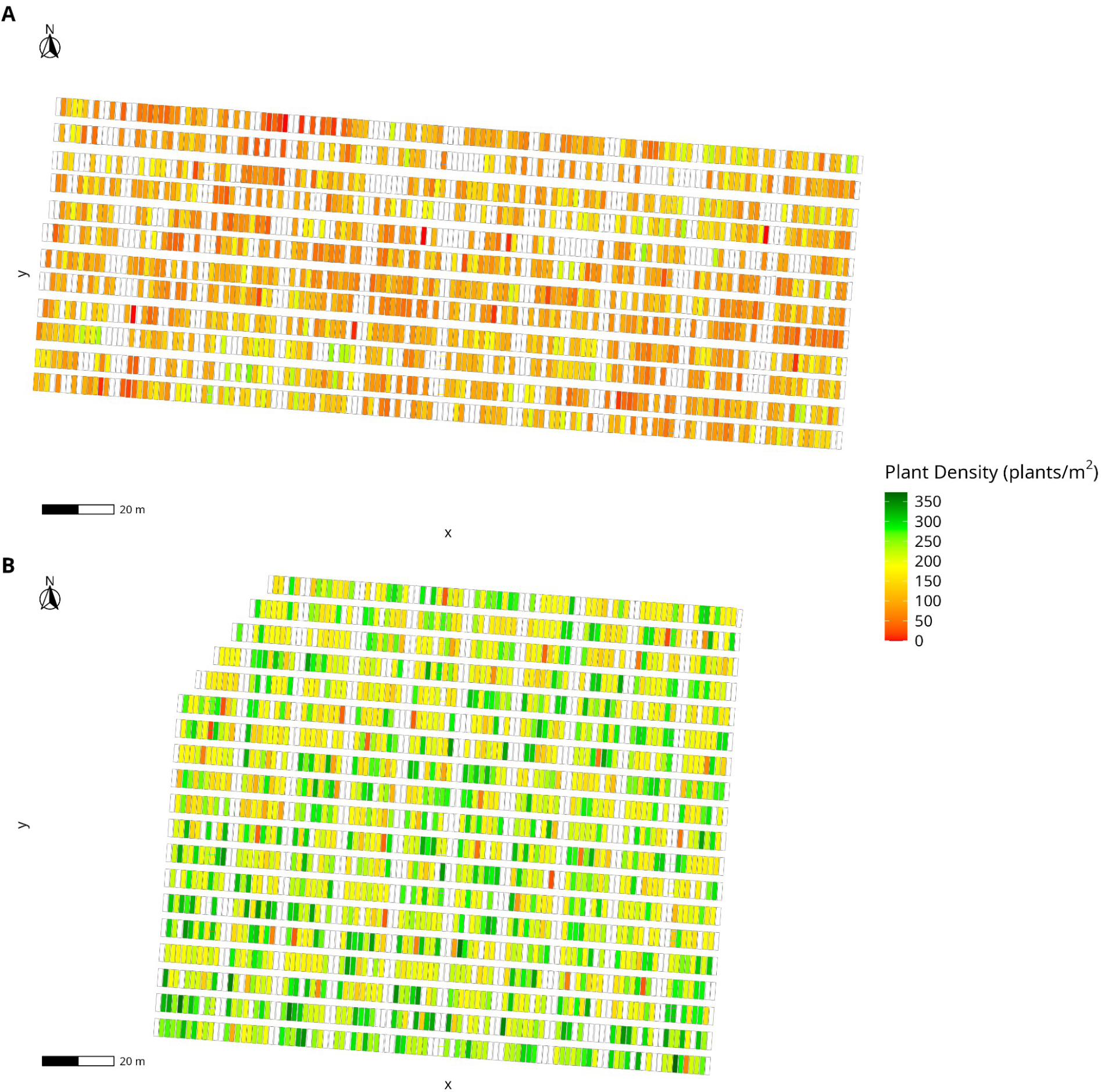
Distribution of predicted plant densities including MBD and CB microplots on the whole platform, 980 microplots in season 1 (A) and 1115 microplots in season 2 (B).

### 3.4 Genetic variation for total plant density

#### 3.4.1 Heritability estimates from manually-scored densities on linear-meters

To validate our methodology, we analysed observed data, i.e. manually-scored plant densities along linear meters of the 40 CB genotypes. We first checked that the mean observed proportion of wheat in IC was a reliable way to infer the wheat plant density from total density in IC (wheat + pea). On both seasons, the corrected wheat plant density in IC aligned with the SC wheat density (supplemental Fig. S5 and supplemental Fig. S6), with high correlation of corrected densities with the true wheat densities, and fairly low RMSE (14.0% and 12.2% for season 1 and 2, respectively). Other correction methods were evaluated, with less accurate results (data not shown).

Then heritability of wheat plant density was first estimated on true wheat densities, using both SC and IC plant densities (i.e. 2 linear meters x 2 blocs x 2 stand types SC/IC for each genotype), with values of 0.61 for season 1 and 0.64 for season 2. Then we estimated heritability from the corrected wheat densities from IC total densities, instead of the true ones, and obtained very similar heritabilities (season 1: 0.61, season 2: 0.69), as expected from the good quality of these corrections.

#### 3.4.2 Heritability estimates from predicted densities

Using the fitted linear model for each season, plant densities were predicted.

Heritability of wheat plant density was then estimated from predicted total plant density of the MBD plots with the same genotypes as CB manual counts (with Flambo as tester for ICWP). Plant density estimates were highly correlated to manual counts, and yielded heritabilities of 0.72 and 0.87 in season 1 and 2, reciprocally, slightly higher than the ones observed on manual counts. When expanding the analysis to the whole platform, i.e. including the Smiley tester (for a total of 980 and 1195 plots in season 1 and 2, reciprocally), we estimated slightly higher heritability for season 1 (0.75) and slightly lower for season 2 (0.85).

## 4. DISCUSSION

### 4.1 Model structure and performance

Our study shows that using a simple linear model, image analysis performed on aerial pictures taken at seedling stage on field trials allows us to predict plant densities with fairly low error rates. Predictions can be performed on both sole or inter-crops, given that this cropping system is provided to the model in training steps. The index of vegetation cover appeared to be the most influential predictor, after log transformation, but integrating other image features improved significantly the quality of predictions.

#### 4.1.1 Fcover is a main, non linear predictor of plant density

Our analyses first highlight that among the different features extracted from aerial pictures taken at seedling stage, images fractional vegetation cover (FCOVER) is the main predictor of plant density in both sole-and inter-crops. The non linearity of this relation is quite unusual, as several previous studies have modelled plant density as a linear function of fractional canopy cover or related vegetation metrics (Jauregui-Besó et al., 2025; Jin et al., 2017; S. Liu et al., 2017; Wilke et al., 2021). (Wilke et al., 2021), for example, explicitly stated that “a higher value of fractional cover at the early seedling stage also indicates a larger number of plants and a linear relation”. To the best of our knowledge, the validity of this assumption has rarely been assessed through an explicit comparison of alternative transformations within a cross-validation framework. Among the formulations evaluated in this study, the logarithmic transformation consistently produced the lowest prediction errors in both growing seasons (supplemental table S1). Although additional environments, species combinations and growth stages would be required to assess the generality of this result, the repeated selection of log(FCOVER) across two contrasted seasons suggests that the non linear relationship between index of vegetation cover and plant density is quite robust.

This non-linearity can be explained by a simple geometric argument: as density increases, the probability that a plant’s projected area overlaps with that of a neighbour plant, rather than bare soil, rises. Preliminary simulations of the geometric footprint of plants from a zenithal view (as in the drone images used), were conducted (data not shown) by modeling wheat leaves as elongated ellipses, taking into account the growth stage, phyllotaxy, and size of the leaves, with plant density along the row varying from 10 to 500 plants.m⁻². These simulations confirmed a non-linear density-to-fcover relationship. However, when considering low densities, a linear relationship is relevant, as expected. It therefore appears that the nature of this relationship depends on the stage of the wheat being studied and the range of seeding densities under consideration: the overlap effect is likely reinforced by high sowing densities, high leaf area by plant, and within-row heterogeneity in seed spacing. These points support three practical considerations: (i) density estimates should be performed at early seedling stage, when overlap between leaves remains limited; (ii) higher accuracy should be observed on field sown with precision seed drills, due to reduced spacing heterogeneity; and (iii) when dealing with high plant densities, underestimation and higher error levels are expected.

#### 4.1.2 Predictors beyond FCOVER improve density estimates

The sequential procedure highlighted the importance of integrating Stand Type (Sole Crop vs Intercrop) and its interaction with FCOVER, which significantly improved the quality of predictions in both seasons. Differences in slopes linking FCOVER and plant density among the different stand types are effectively clear (see supplemental Fig. S4), which can be explained by the difference of plant architecture between wheat and pea, with contrasted leaf surface per plant, requiring adjustments of the model between sole inter-cropping.

Beyond FCOVER and Stand Type, the final models retained a fairly high number of complementary descriptors (24 for season 1 and 15 for season 2, among a total of 76; see Supplemental Table S4 and S6), including canopy cover descriptors, segmentation morphometry, spectral descriptors and texture descriptors. When compared on a common scale (coefficient × standard deviation of the predictor), the number of blobs per linear meter had the largest standardised effect among these complementary descriptors in both seasons, which is consistent with expectations since this index is the one most directly related to plant count. All other descriptors had markedly smaller standardised effects and thus acted as complements rather than as primary predictors. Some overlap was nevertheless found between seasons among these secondary variables, notably spectral (sd_g, sd_ExG, cv_ExG) and morphometric descriptors (mean_blob_area, mean_compactness). The selection of texture predictors shows that they provide complementary information to FCOVER, as highlighted by previous studies (for density: (Shen et al., 2024); for biomass estimates: (Grüner et al., 2021; Yue et al., 2019). Canopy texture features could have captured the Stand-Type factor in the glmnet analyses, excluding it from the predictors. Therefore these features captured other structural variations associated with plant density. Further work is needed to explore which features are the most relevant, and how they improve the prediction accuracy. To improve the accuracy of textural predictors, larger processing windows could be used, accounting for larger spatial neighbourhoods. This would better capture textural information at the plant scale, which consists of patterns made of leaf segments. The GLCM was calculated only on the green channel of the images; it would also be possible to calculate it for each band, as (Zou et al., 2024) did in their study to improve LAI estimation of winter wheat using UAV images.

#### 4.1.3 Model performance

The overall prediction accuracy of selected models was assessed with a statistically sound methodology, i.e. variable selection with nested cross-validation (glmnet), which is not systematically performed. The two final models achieved cross-validation R² values of 0.79 in season 1 (RRMSE = 27.4%) and 0.74 in season 2 (RRMSE = 19.19%). These performances are consistent with those reported for pure wheat stands using RGB or multispectral imagery, although error metrics and validation schemes differ across studies: R² = 0.83 or 0.86 using UAV multispectral imagery ((Wilke et al., 2021) and (Banerjee et al., 2021), respectively), and R² of 0.68-0.90 in RGB imagery (Jauregui-Besó et al., 2025). (S. Liu et al., 2017) achieved higher precision (RRMSE = 12 %), using machine learning on ground-based RGB imagery.

The differences of selected models between the two seasons, both in accuracy and predictor composition, can be linked to the very contrasted meteorological conditions. Season 1 experienced heavy rainfall, lowering germination and emergence rates, and leading to suboptimal and patchy plant densities. Mud deposition on first leaves further decreased growth and also impacted the quality of canopy images. Season 2 benefited from more favourable conditions with uniform emergence and cleaner imagery. The higher predictive quality of the model trained in season 1, is certainly due to the adverse conditions, that globally lowered plant emergence and plant densities (less saturation of FCOVER), but more importantly generated a wide density range, providing a strong statistical signal that favoured higher R² values despite more challenging imagery. Conversely, the more favourable conditions of 2024-2025 produced denser and more uniform stands, reducing density variability and mechanically constraining R², while yielding lower relative prediction errors. These results illustrate that model performance should not be assessed from a single metric and that comparisons across seasons characterised by contrasting levels of biological variability require particular caution.

The present study also addressed a substantially more complex phenotyping context than previous previous studies, as the dataset combined sole crops and wheat-pea intercrops. It also included a large diversity of wheat genotypes. Such diversity in varietal and species composition increases variability in plant architecture and growth, complicating density estimation from canopy imagery. The predictive performances, in the range of reported ones for more homogeneous designs, support the applicability of UAV-based density estimation to breeding-oriented intercrop experiments.

#### 4.1.4 Optimising the training set composition

Working jointly on sole crops and intercrops allowed us to test the importance of the composition of the training set on prediction accuracy. As expected from the significant FCOVER × Stand Type interaction, training-set composition strongly matters, models calibrated exclusively on a single stand type performed poorly when applied to the alternative type, training on wheat sole-crop only being the worst solution. In contrast, all mixed training sets (75/25, 50/50 and 25/75) produced comparable performances on both stand types across the two seasons, almost as good as the natural training set. These results indicate that, once both stand types are present in a trial, integrating both IC and SC provides a major improvement in quality prediction. In our trial, best predictions were achieved with a 25% SC + 75% IC ratio in the number of calibration plots, but this ratio remains flexible, and adjustable to the main objective of the experiment. Therefore a practical recommendation is to integrate manual counts and calibration steps on intercrops in addition to sole crops, as soon as a trial includes both stand types.

### 4.2 Predicting plant density for sole-crop/intercrop breeding

#### 4.2.1 Spatial generalization

The pipeline developed here was successfully applied to 980 and 1195 microplots over two contrasted cropping seasons, offering a practical pathway to non-destructive, high-throughput estimation of plant establishment at the scale of large evaluation trials, such as those used in breeding and varietal evaluation (VCU trials). To the best of our knowledge, no drone imagery has been captured at very low altitudes (less than 10 meters) for high-throughput phenotyping, across the entire scope of a study involving approximately one thousand micro-plots, with a gsd of approximately 0.15 cm. (Koh et al., 2019) monitored approximately 400 microplots, but using drone flights at a higher altitude of 20 meters. Other studies of the same type use similar approaches, namely medium-altitude flights in “photogrammetric” mode, which generally result in centimeter-level resolutions. (Jin et al., 2017) used a hexacopter flying at an altitude between 3 and 7 meters to achieve a very precise GSD of approximately 0.03 cm, for wheat counting, but data collection was limited between 20 to 50 microplots per site due to the time required by the photogrammetric acquisition method used. In their study of barley density, (Wilke et al., 2021) conducted flights at altitudes between 10 and 30 meters, with a GSD of 0.2 cm, but limited to approximately 190 microplots (3 m x 1.4 m). The data acquisition method used in this study consisted of acquisitions at two scales: (i) a single image acquired hovering above micro-parcel at very low altitude, and (ii) a general mosaic of all micro-parcels acquired at medium altitude in photogrammetric mode, which served as a reference for georeferencing the low-altitude image. This method makes it possible to produce very high-resolution maps of a large number of micro-parcels within a reasonable amount of time, and to estimate plant density across ∼1,200 microplots covering approximately 2 ha within ∼6 hours of UAV flight. The manual counts necessary for the calibration were completed within a single day, compared to the several weeks typically required by exhaustive manual counting.

The model, calibrated at the linear-meter scale (∼0.16 m²), predicted realistic plant densities at the microplot scale (∼6.5 m²), within the range of calibration values, thanks to the use of a cross-validation approach, which avoided model overfitting. It is worth emphasizing here the importance of these high-precision manual counts, which provided a reliable ground truth essential to evaluate model accuracy. Among all outliers investigated in the regression between manual counts and plot-scale predictions (Fig. 9), discrepancies could systematically be attributed to within-plot heterogeneity and to non-representative sampling of the counted linear meters. Therefore, using the model at the whole-plot scale improved the prediction of plant density, providing a more robust variable for further analyses.

#### 4.2.2 H2 and breeding

A specificity of the analyzed trial is that it is designed for genetic studies, and presents a high number of wheat genotypes (200). This allowed us to test for a wheat genetic effect on plant density, using an original methodology, i.e. integrating both sole-crop and intercrop plots (doubling the number of plots analysed) to estimate genetic effects on plant density. This strategy consistently improved the estimates, with an unexpected result: high heritability (∼0.8) was estimated for plant density in both trials, when using predicted values at the whole-microplot scale. To our knowledge, this is the first heritability estimate for plant density performed on field trials, even though several studies have documented substantial genetic control over related early-growth traits. Broad-sense heritability estimates of ∼0.86 have been reported for the germination rate measured in Petri dishes on winter wheat panels of the same size to ours (Rajković et al., 2025). More directly relevant to our results, a GWAS conducted on seedling emergence across a global collection of wheat genotypes documented variable broad-sense heritabilities (H² = 0.15 to 0.73; (Francki et al., 2021)). However, these studies were largely conducted under controlled or semi-controlled conditions, and, to our knowledge, no study has estimated the heritability of plant density directly from unselected, large-scale field trials as performed here. As highlighted by (Francki et al., 2021) heritability measures of plant density also integrate maternal effects: growth conditions can impact seed viability and vigor (seed size), while poor conservation conditions can degrade germination rate, either inflating or decreasing heritability estimates. This source of variation, distinct from the genetic effect of the evaluated genotype itself, should be kept in mind when interpreting heritability estimates for plant density and related early-growth traits. More broadly, plant density is only one of several early-growth traits for which genetic variation has been documented in wheat and associated with agronomic performance. (Rebetzke & Richards, 1999) reported high broad-sense heritability for early vigour (H² = 87%), together with strong positive genetic correlations with leaf breadth and length. Genetic variation in specific leaf area itself has since been shown to offer a promising and largely untapped lever for improving early vigour (Rebetzke et al., 2004). Coleoptile length, a trait directly relevant to seedling emergence and therefore connected to the plant density estimates presented here, has similarly been shown to combine high heritability (H² = 0.60 to 0.86) with exploitable genetic variation across breeding populations (Murphy et al., 2008). Taken together, these results suggest that plant density could be usefully integrated within this broader complex of early-vigour traits, and that the UAV-based phenotyping approach developed here may offer a scalable route to jointly characterise several of them in field conditions.

Beyond this broader trait perspective, methodological refinements could also strengthen the robustness of the heritability estimates presented here. A useful development would be to integrate the prediction uncertainty from the glmnet regression (using bootstrap or conformal prediction) to estimate heritabilities with associated confidence intervals.

### 4.3 Limits to the method and perspectives

When developing breeding programmes targeting wheat-legume intercrops (Moutier et al., 2022; Sampoux et al., 2020), phenotyping both stand types with the same pipeline becomes a methodological requirement (Salomon et al., 2026). Studies explicitly targeting cereal-legume intercrops at the scale of a multi-genotype evaluation trial remain scarce. Among the few studies addressing UAV-based density estimation in pluri-specific canopies, (Koh et al., 2019) focused on early mixed growth stages in safflower, and (Vong et al., 2021) compared three crop types in maize. Our study extends here UAV-based density estimation to a wheat-pea intercrop system involving 200 wheat genotypes, with sole-crop and intercrop plots evaluated under a unified pipeline. As the method is centered on vegetation index, early measurements should be favoured, as they minimise the confounding contribution of individual plant growth to the signal; conversely, later measurements increasingly reflect early vigour rather than density per se. The sequential analysis of early growth stages can be a way to better integrate plant size variations due to differences in emergence timing and genotypes. Beyond its use as a phenotypic trait in its own right, density prediction is also relevant to the broader analysis of agronomic and genetic field trials. UAV-derived indices have already been shown to improve the fit of spatial models in wheat breeding yield trials, helping to disentangle genetic from environmental sources of variation (Herr et al., 2024); predicted plant density could similarly serve as a covariate to characterise and correct for establishment-related spatial heterogeneity. Altogether, this pipeline could substantially reduce field workload at the beginning of the season, while providing an early indicator of crop establishment that allows breeders to anticipate within-trial density variability and adjust downstream analyses accordingly.

For image-based approaches, it is very important to pay close attention to the quality of the acquired data, as this quality largely determines the results obtained. Compared to deep-learning-based approaches, the pipeline presented here requires recalibration each season to account for changes in crop growth stage and image quality, a clear limitation relative to a well-trained neural network. In return, our approach does not depend on large annotated training datasets, remains interpretable, and readily discriminates between species based on canopy architecture. Future comparisons between the two approaches should also weigh the possibility, offered here, of easily integrating additional indices to characterise intra-microplot heterogeneity, an aspect less straightforward to extract from deep-learning models. A last critical improvement will be to integrate the discrimination of the intercropped species (Munz & Reiser, 2020; Pranga et al., 2024), to achieve species-specific density estimates.

## AUTHOR CONTRIBUTIONS

**Virgilio Freitas:** Investigation; data curation; formal analysis; methodology; software; writing original draft

**Jérôme Enjalbert**: Investigation; formal analysis; methodology; resources; supervision; validation;writing original draft

**Timothée Flutre**: Investigation; formal analysis; methodology; software; resources; supervision; validation; writing original draft

**Jean-Marc Gilliot**: Investigation; formal analysis; methodology; software; resources; supervision; validation; writing original draft

## ACKNOWLEDGMENTS

We would like to thank the INRAE unit Versailles-Saclay (UEVS) for its management of the trials at the “Le Moulon” experimental farm.

This work, conducted as part of the coBreeding project (Co-design of animal and plant breeding schemes with multi-performance objectives), received government funding, notably the PhD grant of V. Freitas, administered by the French National Research Agency (ANR) under the France 2030 program, grant number ANR-22-PEAE-0003. The field experiment was carried-out within the MoBiDiv project, administered by ANR under the PPR “Cultiver Protéger Autrement” program (ANR-20-PCPA-0006).

## DATA AVAILABILITY STATEMENT

The data and scripts supporting the findings of this study are available from the corresponding author upon reasonable request.

## Abbreviations

AGL: above ground level
CB: cobreeding
CV: cross validation
DEM: digital elevation model
Fcover: fractional vegetation cover
GCP: ground control point
GLCM: grey-level co-occurrence matrix
GMC: grain moisture content
GSD: ground sample distance
GW: grain weight
GY: grain yield
IC: intercrop
ICWP: intercrop wheat pea
LER: land equivalent ratio
MBD: MoBiDiv
OBIA: object-based image analysis
PC: plant count
PD: plant density
PLSR: partial least squares regression
RGB: red green blue
RMSE: root mean square error
RRMSE: relative root mean square error
RTK: real-time kinematic
SC: sole crop
SCP: sole crop pea
SCW: sole crop wheat
SVM: support vector machine
totGY: total grain yield
UAV: uncrewed aerial vehicle
UEVS: Versailles-Saclay experimental unit.

## SUPPLEMENTAL MATERIAL

**Supplemental Fig. S1:**
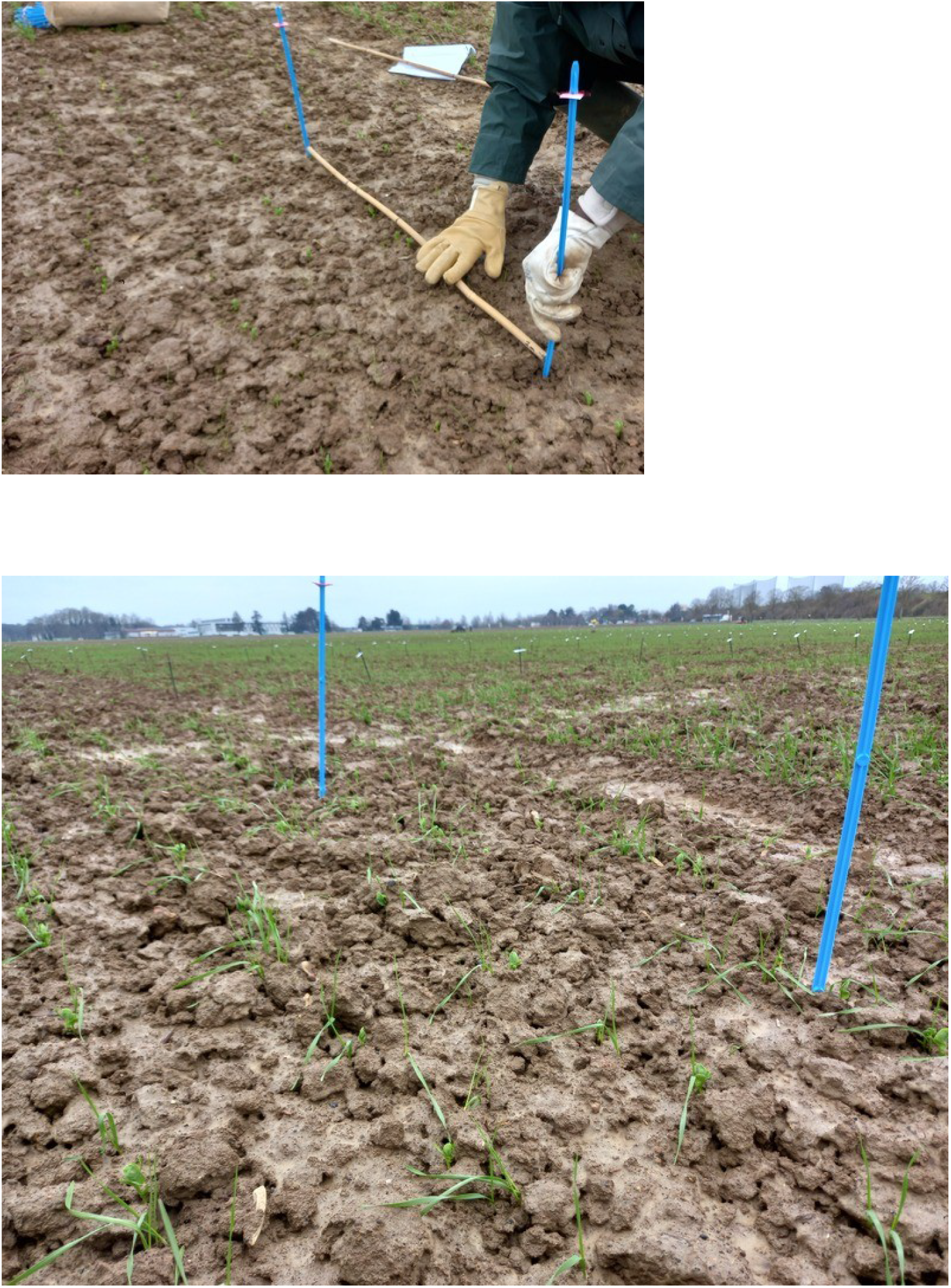
Installation of the linear meters.

**Supplemental Fig. S2:**
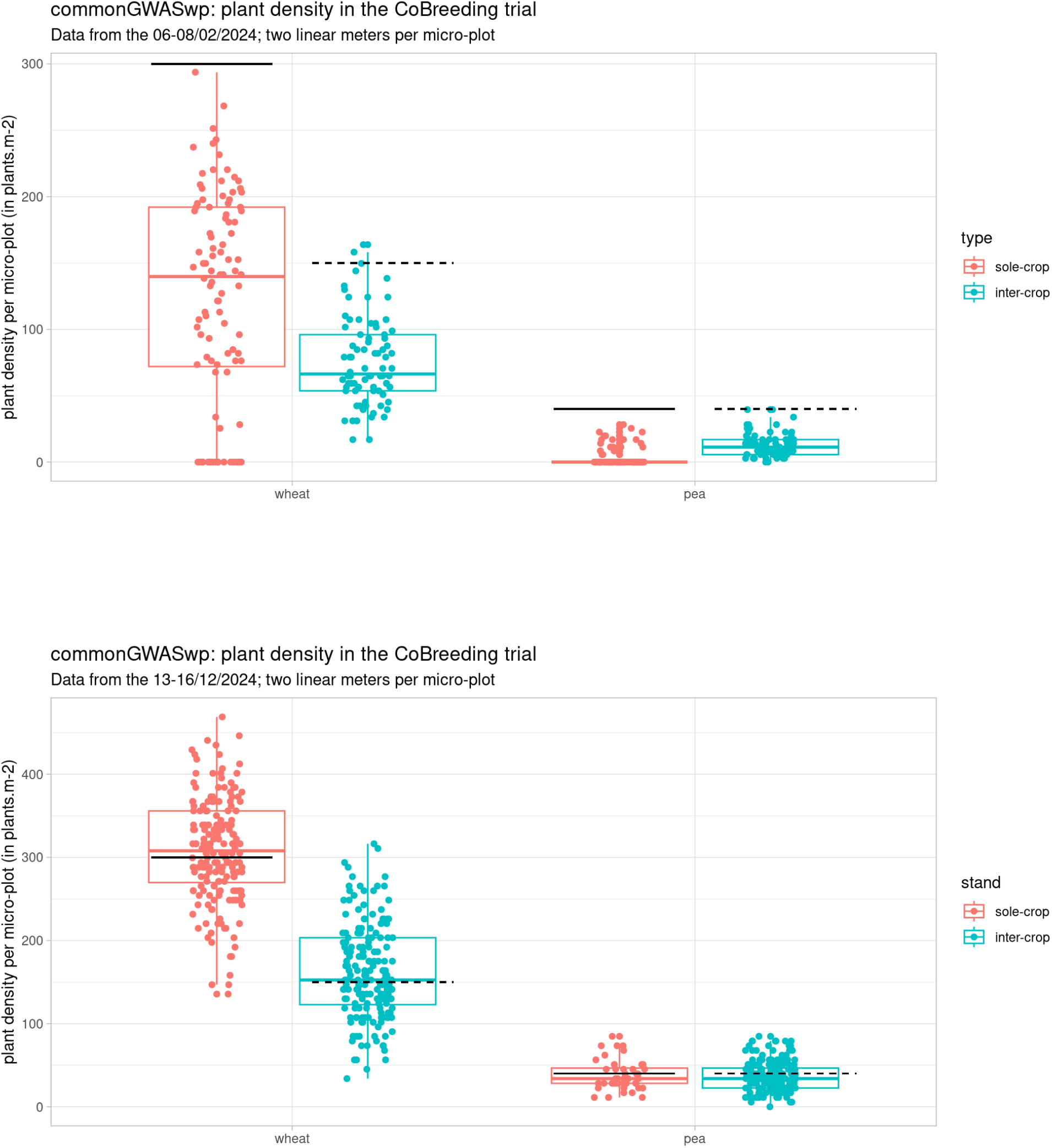
Plant density per species by manual scoring along the linear meters (one point per linear meters)

**Supplemental Fig. S3:**
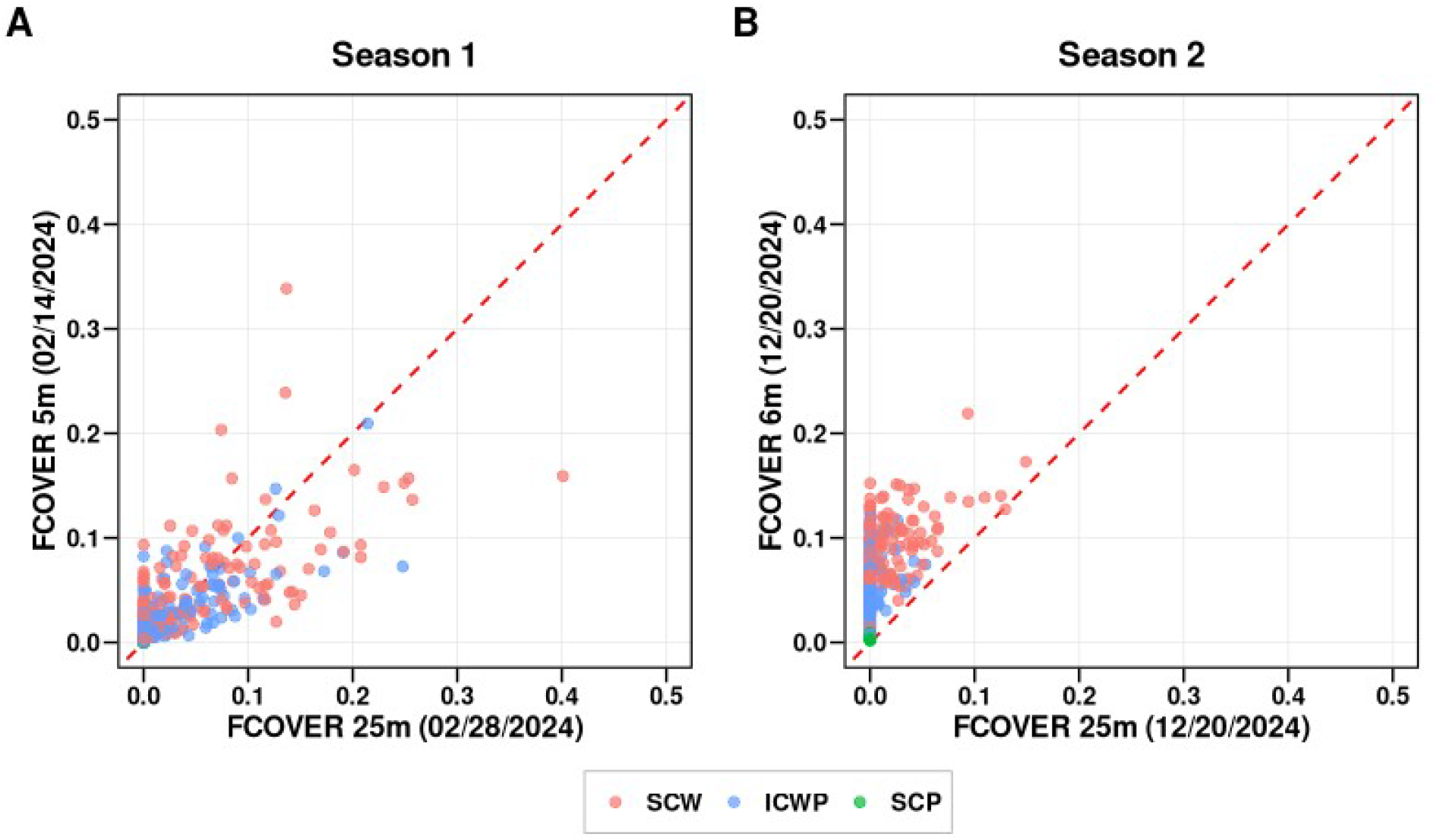
Relationship between FCOVER of each micro-plot derived from RGB images from UAV flights at high (25 meters) versus low (5 or 6 meters) altitudes, in season 1 (A) and season 2 (B).

**Supplemental Fig. S4:**
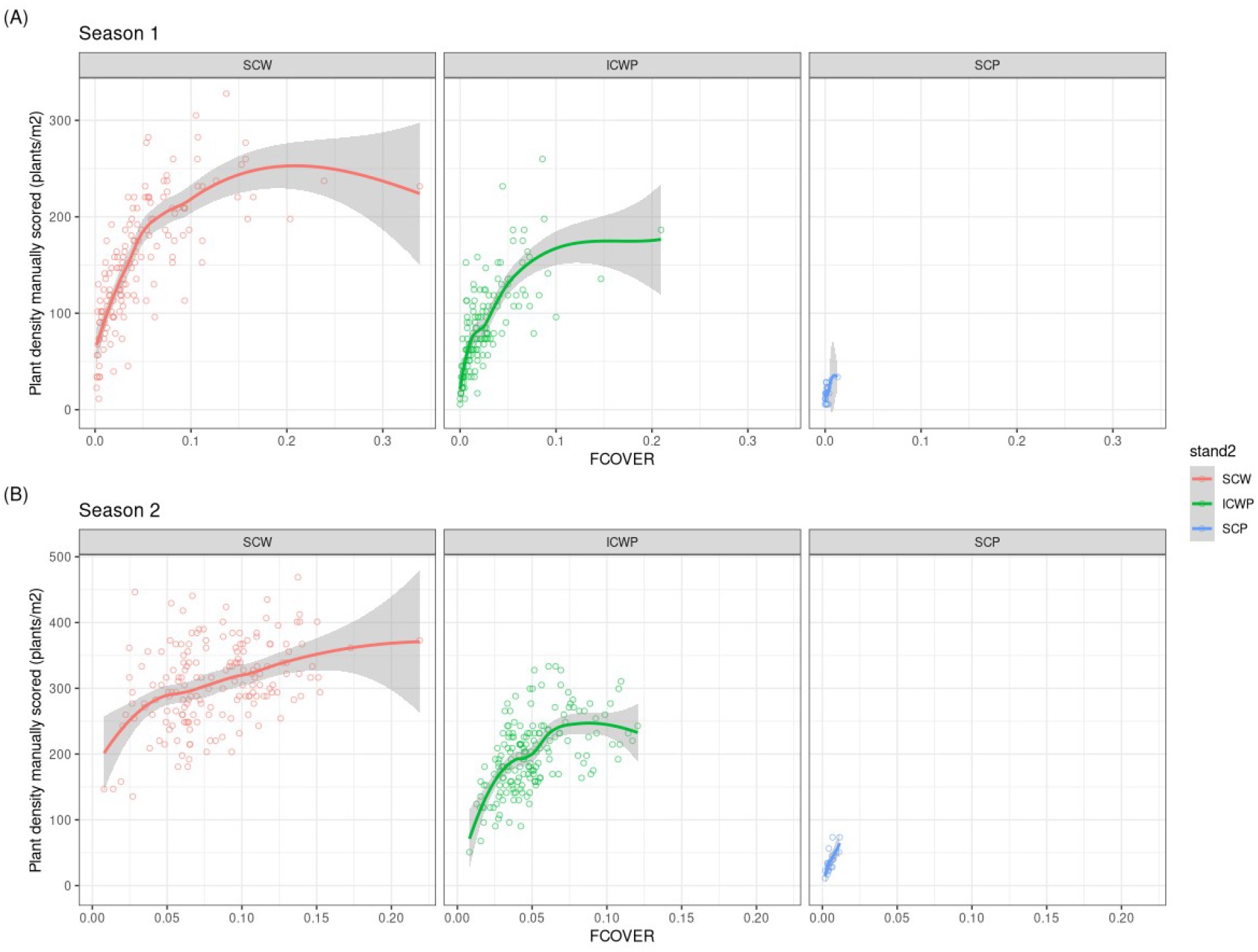
Relation between manually-scored plant density and FCOVER

**Supplemental Fig. S5:**
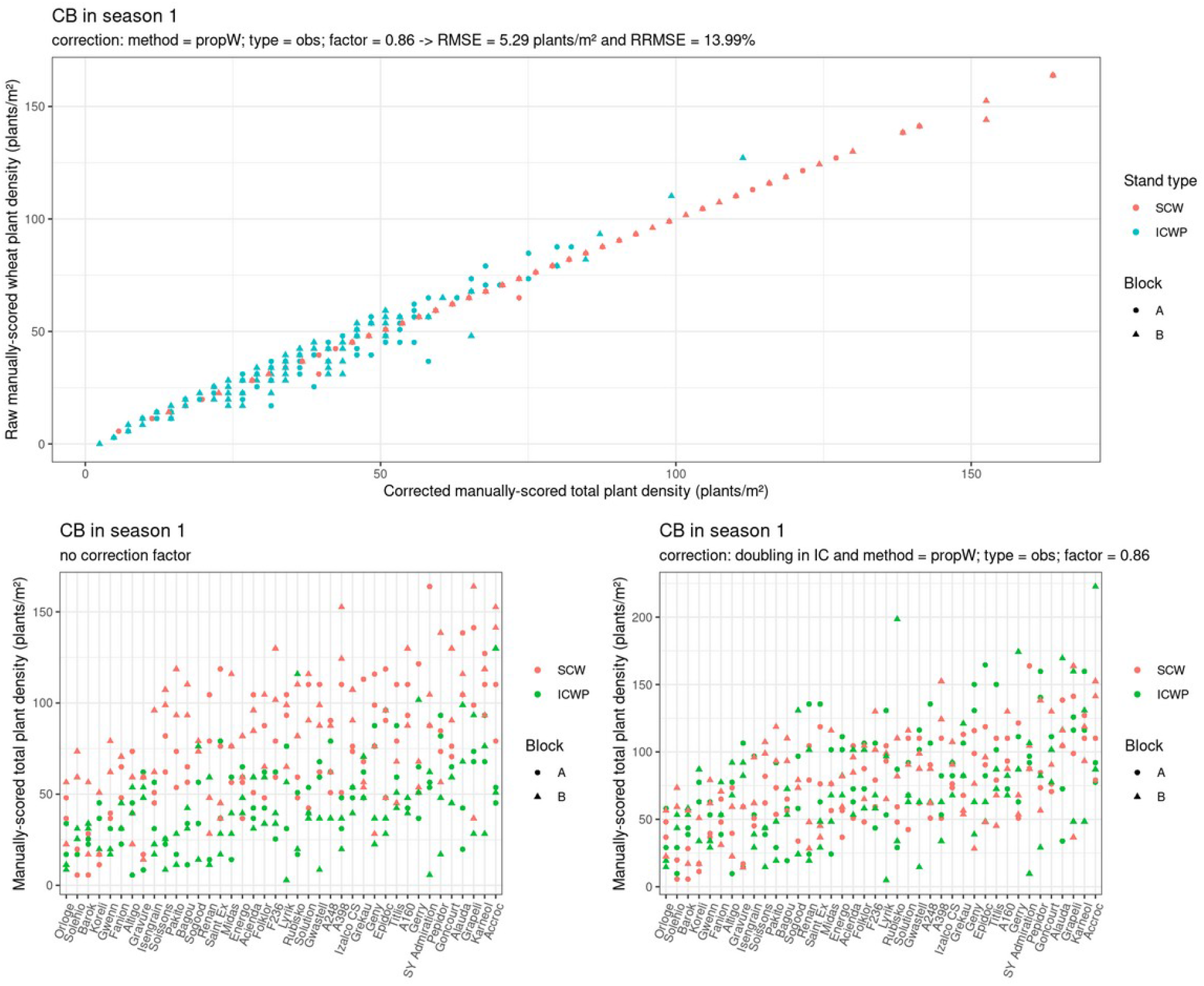
Raw wheat versus corrected total manually-scored plant densities from CoBreeding (CB) micro-plots in season 1

**Supplemental Fig. S6:**
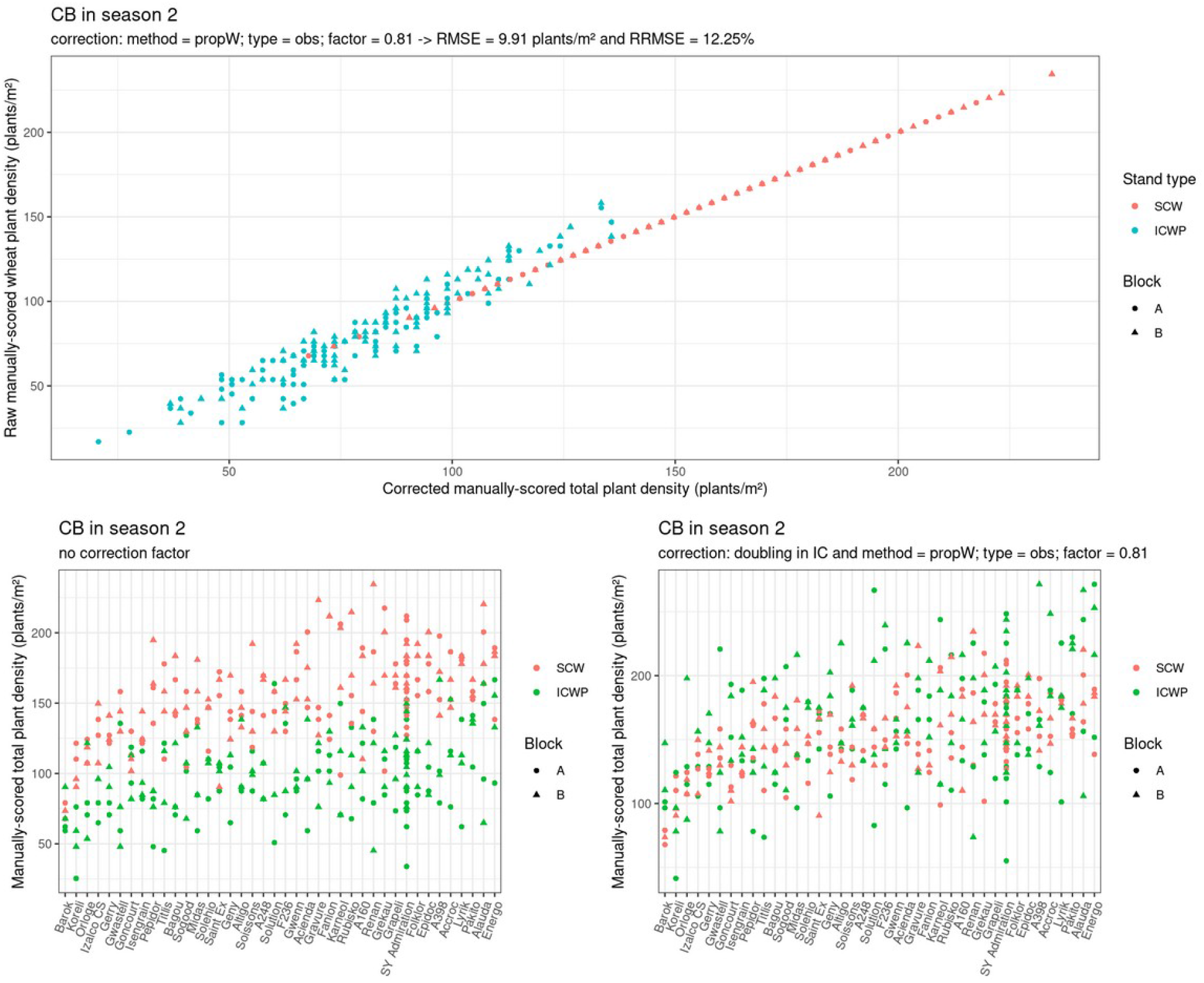
Raw wheat versus corrected total manually-scored plant densities from CoBreeding (CB) micro-plots in season 2

---

**Supplemental Table S1:** Investigation of FCOVER transformations, cross-validated performance of plant density prediction using Fcover transformation as predictor.

| Fcover input | Season 1 |  |  | Season 2 |  |  |
| --- | --- | --- | --- | --- | --- | --- |
|  | R<sup>2</sup> | RMSE | RRMSE | R<sup>2</sup> | RMSE | RRMSE |
|  | — | g m<sup>-2</sup> | % | — | g m<sup>-2</sup> | % |
| Fcover (raw) | 0.448 | 50.79 | 44.6 | 0.45 | 68.50 | 28.4 |
| sqrt(Fcover) | 0.629 | 41.63 | 36.5 | 0.53 | 63.40 | 26.3 |
| log(Fcover) | 0.641 | 40.95 | 35.9 | 0.56 | 61.34 | 25.4 |
| Fcover<sup>2</sup> | -0.027 | 69.22 | 60.7 | 0.296 | 77.58 | 32.2 |
| Fcover<sup>3</sup> | -0.411 | 80.82 | 70.9 | 0.144 | 85.59 | 35.5 |

**Supplemental Table S2:** Investigation of crop type, effect of including “type” variable on cross validated plant density prediction model, using the retained transformation of fcover, for season 1 and 2.

| Model | Season 1 |  |  | Season 2 |  |  |
| --- | --- | --- | --- | --- | --- | --- |
|  | R<sup>2</sup> | RMSE | RRMSE | R<sup>2</sup> | RMSE | RRMSE |
|  | — | g m<sup>-2</sup> | % | — | g m<sup>-2</sup> | % |
| M1 | 0.623 | 42.01 | 36.9 | 0.559 | 61.42 | 25.5 |
| M2 | 0.707 | 37.02 | 32.5 | 0.666 | 53.48 | 22.2 |
| M3 | 0.724 | 35.89 | 31.5 | 0.668 | 53.26 | 22.1 |

**Supplemental Table S3:** Elastic-net hyperparameter tuning grid for the plant density prediction model: cross-validated RMSE and number of non-zero coefficients across α values, season 1.

| $\alpha$ | $\lambda(1se)$ | CV_RMSE | nzero |
| --- | --- | --- | --- |
| 0.000 | 61.946 | 31.328 | 79 |
| 0.001 | 61.946 | 31.343 | 63 |
| 0.008 | 14.85 | 30.540 | 53 |
| 0.027 | 1.904 | 30.332 | 56 |
| 0.064 | 1.404 | 30.313 | 51 |
| 0.125 | 1.144 | 30.317 | 40 |
| 0.216 | 0.798 | 30.268 | 36 |
| 0.343 | 0.605 | 30.267 | 33 |
| 0.512 | 0.405 | 30.192 | 34 |
| 0.729 | 0.284 | 30.139 | 32 |
| 1<sup>a</sup> | 0.207 | 30.092 | 27 |

**Supplemental Table S4:** Variables retained by the elastic-net model and their coefficients, season 1.

| variable | coef_raw | sd_x | coef_std |
| --- | --- | --- | --- |
| typepure | 91.427 | 0.499 | 45.674 |
| FCOVER_log:typepure | 15.035 | 2.094 | 31.495 |
| sd_g | 6962.794 | 0.003 | 25.761 |
| FCOVER_log | 16.507 | 1.193 | 19.695 |
| n_blobs_perM | 0.333 | 53.301 | 17.773 |
| frac_007 | -220.220 | 0.064 | -14.115 |
| frac_009 | 348.076 | 0.038 | 13.392 |
| glcm_contrast_w11_a45 | -10.970 | 1.165 | -12.788 |
| mean_ExG | -948.800 | 0.013 | -12.4135 |
| glcm_homogeneity_w11_a0 | -72.410 | 0.076 | -5.526 |
| frac_003 | -20.943 | 0.262 | -5.493 |
| glcm_contrast_w11_a90 | -4.827 | 0.730 | -3.526 |
| mean_blob_area | -10336.028 | 0.0002 | -3.088 |
| frac_005 | 19.900 | 0.151 | 3.005 |
| mean_compactness | -9.704 | 0.273 | -2.654 |
| sd_VARI | -2.469 | 1.041 | -2.572 |
| glcm_contrast_w11_a0 | 3.494 | 0.733 | 2.563 |
| glcm_homogeneity_w15_a0 | -19.170 | 0.076 | -1.463 |
| glcm_contrast_w7_a45 | -1.0009 | 1.166 | -1.167 |
| glcm_contrast_w5_a90 | -1.506 | 0.730 | -1.101 |
| cv_ExG | -3.074 | 0.350 | -1.077 |
| glcm_contrast_w5_a0 | 0.806 | 0.734 | 0.591 |
| glcm_contrast_w9_a45 | -0.248 | 1.165 | -0.289 |
| glcm_contrast_w5_a45 | -0.007 | 1.166 | -0.008 |
| glcm_contrast_w9_a0 | 0.010 | 0.733 | 0.007 |
| glcm_contrast_w7_a0 | 0.007 | 0.733 | 0.005 |
| glcm_contrast_w15_a0 | 0.002 | 0.733 | 0.001 |

**Supplemental Table S5:** Elastic-net hyperparameter tuning grid for the plant density prediction model: cross-validated RMSE and number of non-zero coefficients across α values, season 2.

| $\alpha$ | $\lambda(1se)$ | CV_RMSE | nzero |
| --- | --- | --- | --- |
| 0 | 59.170 | 45.565 | 79 |
| 0.001 | 53.914 | 45.471 | 77 |
| 0.008 | 47.544 | 45.557 | 56 |
| 0.027 | 33.716 | 45.574 | 33 |
| 0.064 | 23.992 | 45.561 | 21 |
| 0.125 | 14.796 | 45.477 | 19 |
| 0.216<sup>a</sup> | 9.397 | 45.438 | 18 |
| 0.343 | 6.494 | 45.494 | 17 |
| 0.512 | 4.775 | 45.575 | 15 |
| 0.729 | 3.353 | 45.527 | 15 |
| 1 | 2.444 | 45.502 | 12 |

**Supplemental Table S6:**
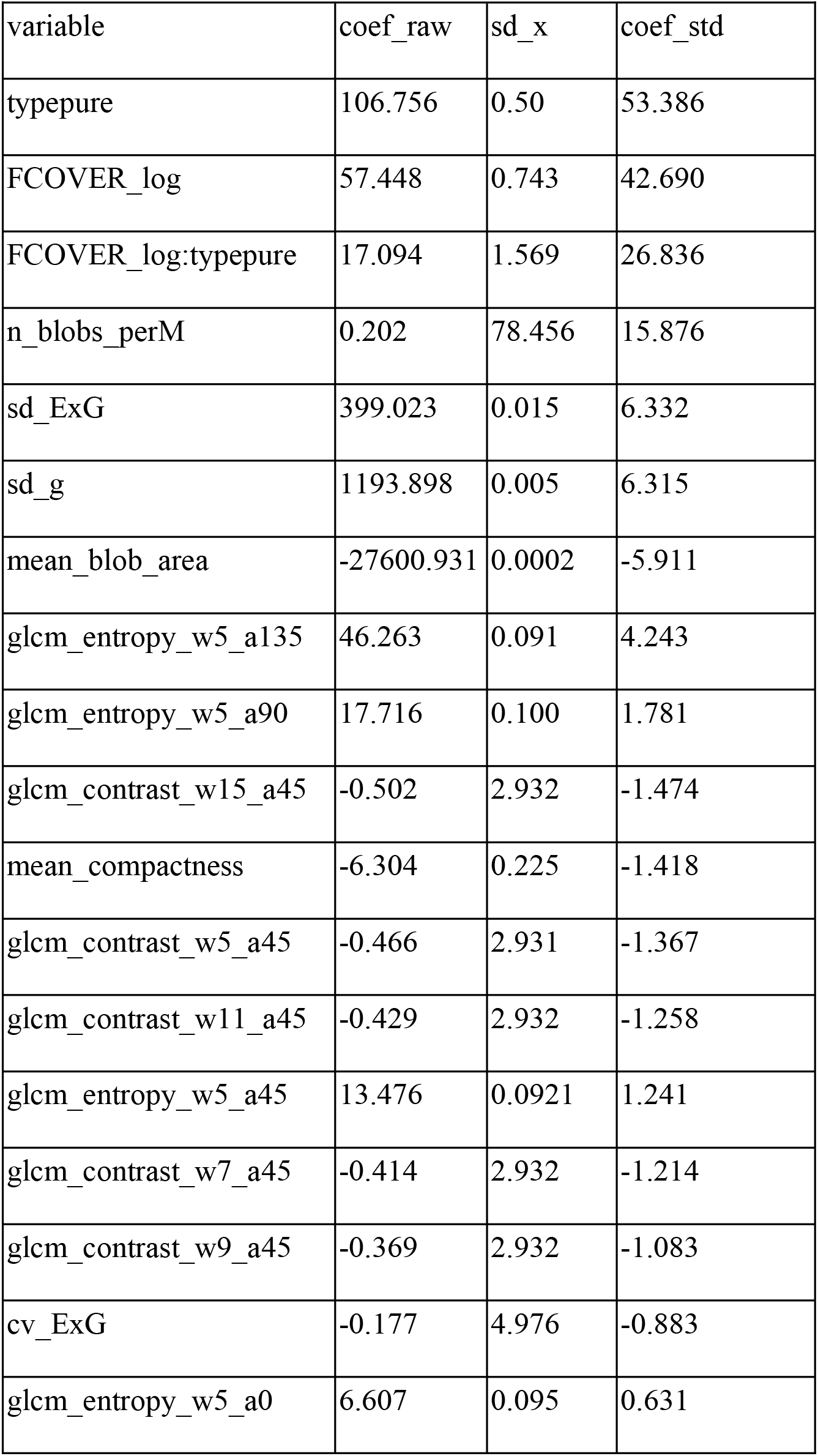
Variables retained by the elastic-net model and their coefficients, season 2.

| variable | coef_raw | sd_x | coef_std |
| --- | --- | --- | --- |
| typepure | 106.756 | 0.50 | 53.386 |
| FCOVER_log | 57.448 | 0.743 | 42.690 |
| FCOVER_log:typepure | 17.094 | 1.569 | 26.836 |
| n_blobs_perM | 0.202 | 78.456 | 15.876 |
| sd_ExG | 399.023 | 0.015 | 6.332 |
| sd_g | 1193.898 | 0.005 | 6.315 |
| mean_blob_area | -27600.931 | 0.0002 | -5.911 |
| glcm_entropy_w5_a135 | 46.263 | 0.091 | 4.243 |
| glcm_entropy_w5_a90 | 17.716 | 0.100 | 1.781 |
| glcm_contrast_w15_a45 | -0.502 | 2.932 | -1.474 |
| mean_compactness | -6.304 | 0.225 | -1.418 |
| glcm_contrast_w5_a45 | -0.466 | 2.931 | -1.367 |
| glcm_contrast_w11_a45 | -0.429 | 2.932 | -1.258 |
| glcm_entropy_w5_a45 | 13.476 | 0.0921 | 1.241 |
| glcm_contrast_w7_a45 | -0.414 | 2.932 | -1.214 |
| glcm_contrast_w9_a45 | -0.369 | 2.932 | -1.083 |
| cv_ExG | -0.177 | 4.976 | -0.883 |
| glcm_entropy_w5_a0 | 6.607 | 0.095 | 0.631 |

**Supplemental Table S7:** Training and validation set composition for evaluating the effect of training-set composition on plant density prediction.

| Train | Training set ($n_t=120$) | Target | Validation set ($n_v=40$) | Test type |
| --- | --- | --- | --- | --- |
| 100SC | 120 SC among 160 SC (R times) | SC | remaining 40 SC | cross validation |
|  |  | IC | 40 IC sampled among 160 IC (R times) | test validation |
| 100IC | 120 IC among 160 IC (R times) | IC | remaining 40 IC | cross validation |
|  |  | SC | 40 SC sampled among 160 SC (R times) | test validation |
| 75IC-25SC | 90 IC among 160 IC + 30 SC among 160 SC (R times) | SC | 40 SC sampled among remaining 130 SC (R times) | test validation |
|  |  | IC | 40 IC sampled among remaining 70 IC (R times) | test validation |
| 50SC-50IC | 80 IC among 160 IC + 80 SC among 160 SC (R times) | SC | 40 SC sampled among remaining 80 SC (R times) | test validation |
|  |  | IC | 40 IC sampled among remaining 80 SC (R times) | test validation |
| 25SC-25IC | 30 IC among 160 IC + 90 SC among 160 SC (R times) | SC | 40 SC sampled among remaining 70 SC (R times) | test validation |
|  |  | IC | 40 IC sampled among remaining 130 SC (R times) | test validation |

## REFERENCES

1. Aich, S., Josuttes, A., Ovsyannikov, I., Strueby, K., Ahmed, I., Duddu, H. S., Pozniak, C., Shirtliffe, S., & Stavness, I. (2018). DeepWheat: Estimating Phenotypic Traits from Crop Images with Deep Learning. 2018 IEEE Winter Conference on Applications of Computer Vision (WACV), 323–332. 10.1109/WACV.2018.00042

2. Araus, J. L., & Cairns, J. E. (2014). Field high-throughput phenotyping: The new crop breeding frontier. Trends in Plant Science. 10.1016/j.tplants.2013.09.008

3. Bančič, J., Gorjanc, G., & Tolhurst, D. J. (2024). A framework for simulating genotype-by-environment interaction using multiplicative models. TAG. Theoretical and Applied Genetics. Theoretische Und Angewandte Genetik, 137(8), 197. 10.1007/s00122-024-04644-7

4. Banerjee, B., Sharma, V., Spangenberg, G., & Kant, S. (2021). Machine Learning Regression Analysis for Estimation of Crop Emergence Using Multispectral UAV Imagery. Remote Sensing, 13, 2918. 10.3390/rs13152918

5. Bates, D., Mächler, M., Bolker, B., & Walker, S. (2015). Fitting Linear Mixed-Effects Models Using lme4. Journal of Statistical Software, 67, 1–48. 10.18637/jss.v067.i01

6. Bedoussac, L., Journet, E.-P., Hauggaard-Nielsen, H., Naudin, C., Corre-Hellou, G., Jensen, E. S., Prieur, L., & Justes, E. (2015). Ecological principles underlying the increase of productivity achieved by cereal-grain legume intercrops in organic farming. A review. Agronomy for Sustainable Development, 35(3), 911–935. 10.1007/s13593-014-0277-7

7. Beillouin, D., Ben-Ari, T., Malézieux, E., Seufert, V., & Makowski, D. (2021). Positive but variable effects of crop diversification on biodiversity and ecosystem services. Global Change Biology, 27(19), 4697–4710. 10.1111/gcb.15747

8. Demie, D. T., Döring, T. F., Finckh, M. R., van der Werf, W., Enjalbert, J., & Seidel, S. J. (2022). Mixture × Genotype Effects in Cereal/Legume Intercropping. Frontiers in Plant Science, 13, 846720. gaudio

9. Dunnington, D., Vanderhaeghe, F., Caha, J., & Muenchow, J. (2024). R package qgisprocess: Use QGIS processing algorithms. 10.5281/zenodo.10631683

10. Francki, M. G., Stainer, G. S., Walker, E., Rebetzke, G. J., Stefanova, K. T., & French, R. J. (2021). Phenotypic Evaluation and Genetic Analysis of Seedling Emergence in a Global Collection of Wheat Genotypes (Triticum aestivum L.) Under Limited Water Availability. Frontiers in Plant Science, 12. 10.3389/fpls.2021.796176

11. Friedman, J. H., Hastie, T., & Tibshirani, R. (2010). Regularization Paths for Generalized Linear Models via Coordinate Descent. Journal of Statistical Software, 33, 1–22. 10.18637/jss.v033.i01

12. Gano, B., Bhadra, S., Vilbig, J. M., Ahmed, N., Sagan, V., & Shakoor, N. (2024). Drone-based imaging sensors, techniques, and applications in plant phenotyping for crop breeding: A comprehensive review. The Plant Phenome Journal, 7(1), e20100. 10.1002/ppj2.20100

13. Gnädinger, F., & Schmidhalter, U. (2017). Digital Counts of Maize Plants by Unmanned Aerial Vehicles (UAVs). Remote Sensing, 9(6), 544. 10.3390/rs9060544

14. Grüner, E., Astor, T., & Wachendorf, M. (2021). Prediction of Biomass and N Fixation of Legume–Grass Mixtures Using Sensor Fusion. Frontiers in Plant Science, 11. 10.3389/fpls.2020.603921

15. Haralick, R. M., Shanmugam, K., & Dinstein, I. (1973). Textural Features for Image Classification. *IEEE Transactions on Systems, Man, and Cybernetics*, SMC*-*3(6), 610–621. 10.1109/TSMC.1973.4309314

16. Haug, B., Messmer, M. M., Enjalbert, J., Goldringer, I., Flutre, T., Mary-Huard, T., & Hohmann, P. (2023). New insights towards breeding for mixed cropping of spring pea and barley to increase yield and yield stability. Field Crops Research, 297, 108923. 10.1016/j.fcr.2023.108923

17. Haug, B., Messmer, M. M., Enjalbert, J., Goldringer, I., Forst, E., Flutre, T., Mary-Huard, T., & Hohmann, P. (2021). Advances in Breeding for Mixed Cropping – Incomplete Factorials and the Producer/Associate Concept. Frontiers in Plant Science, 11, 620400. 10.3389/fpls.2020.620400

18. Herr, A. W., Campbell, K. G., Li, X., & Carter, A. H. (2024). Spatial analysis with unoccupied aircraft systems data in wheat breeding yield trials. The Plant Phenome Journal, 7(1), e70007. 10.1002/ppj2.70007

19. Jauregui-Besó, J., Gracia-Romero, A., Carrera, C. S., Lopes, M. da S., Araus, J. L., & Kefauver, S. C. (2025). Winter wheat plant density determination: Robust predictions across varied agronomic conditions using multiscale RGB imaging. Smart Agricultural Technology, 11, 100921. 10.1016/j.atech.2025.100921

20. Jensen, E. S., Carlsson, G., & Hauggaard-Nielsen, H. (2020). Intercropping of grain legumes and cereals improves the use of soil N resources and reduces the requirement for synthetic fertilizer N: A global-scale analysis. Agronomy for Sustainable Development, 40(1), 5. 10.1007/s13593-020-0607-x

21. Jensen, S. M., Jørgensen, J. R., & Rasmussen, J. (2023). A non-replicated experimental design in faba bean (Vicia faba L.) variety testing – Embracing plant density variation as a strength. European Journal of Agronomy, 148, 126851. 10.1016/j.eja.2023.126851

22. Jin, X., Liu, S., Baret, F., Hemerlé, M., & Comar, A. (2017). Estimates of plant density of wheat crops at emergence from very low altitude UAV imagery. Remote Sensing of Environment, 198, 105–114. 10.1016/j.rse.2017.06.007

23. Jin, X., Zarco-Tejada, P. J., Schmidhalter, U., Reynolds, M. P., & Et., A. (2021). High-Throughput Estimation of Crop Traits: A Review of Ground and Aerial Phenotyping Platforms. IEEE Geoscience and Remote Sensing Magazine. 10.1109/MGRS.2020.2998816

24. Koh, J. C. O., Hayden, M., Daetwyler, H., & Kant, S. (2019). Estimation of crop plant density at early mixed growth stages using UAV imagery. Plant Methods. 10.1186/s13007-019-0449-1

25. Liu, S., Baret, F., Andrieu, B., Burger, P., & Hemmerlé, M. (2017). Estimation of Wheat Plant Density at Early Stages Using High Resolution Imagery. Frontiers in Plant Science, 8, 739. 10.3389/fpls.2017.00739

26. Liu, Y., Feng, H., Yue, J., Jin, X., Li, Z., & Yang, G. (2022). Estimation of potato above-ground biomass based on unmanned aerial vehicle red-green-blue images with different texture features and crop height. Frontiers in Plant Science, 13. 10.3389/fpls.2022.938216

27. Lu, J., Qi, Q., Zheng, G., Eitel, J. U. H., Zhang, Q., Zhang, J., Chen, S., Zhang, F., Fang, W., Guan, Z., & Chen, F. (2025). High-Throughput Field Phenotyping Using Unmanned Aerial Vehicles (UAVs) for Rapid Estimation of Photosynthetic Traits. Plant Phenomics, 7(2), 100045. 10.1016/j.plaphe.2025.100045

28. Moutier, N., Baranger, A., Fall, S., Hanocq, E., Marget, P., Floriot, M., & Gauffreteau, A. (2022). Mixing Ability of Intercropped Wheat Varieties: Stability Across Environments and Tester Legume Species. Frontiers in Plant Science, 13, 877791. 10.3389/fpls.2022.877791

29. Munz, S., & Reiser, D. (2020). Approach for Image-Based Semantic Segmentation of Canopy Cover in Pea–Oat Intercropping. Agriculture, 10(8), Article 8. 10.3390/agriculture10080354

30. Murphy, K., Balow, K., Lyon, S. R., & Jones, S. S. (2008). Response to selection, combining ability and heritability of coleoptile length in winter wheat. Euphytica, 164(3), 709–718. 10.1007/s10681-008-9692-7

31. Nuijten, R. J. G., Kooistra, L., & De Deyn, G. B. (2019). Using Unmanned Aerial Systems (UAS) and Object-Based Image Analysis (OBIA) for Measuring Plant-Soil Feedback Effects on Crop Productivity. Drones, 3(3), 54. 10.3390/drones3030054

32. Olson, N. A., Trostle, C., Meyer, R., & Hulke, B. S. (2024). Canopy closure, yield, and quality under heterogeneous plant spacing in sunflower. Agronomy Journal, 116(5), 2275–2283. 10.1002/agj2.21655

33. Osco, L., Junior, J., Ramos, A. P., Jorge, L., Fatholahi, S. N., Silva, J., Matsubara, E., Pistori, H., Gonçalves, W., & Li, J. (2021). A Review on Deep Learning in UAV Remote Sensing. International Journal of Applied Earth Observation and Geoinformation, 102, 102456. 10.1016/j.jag.2021.102456

34. Otsu, N. (1979). A Threshold Selection Method from Gray-Level Histograms. *IEEE Transactions on Systems*, Man, and Cybernetics, 9(1), 62–66. 10.1109/TSMC.1979.4310076

35. Pathak, H., Igathinathane, C., Zhang, Z., Archer, D., & Hendrickson, J. (2022). A review of unmanned aerial vehicle-based methods for plant stand count evaluation in row crops. Computers and Electronics in Agriculture, 198, 107064. 10.1016/j.compag.2022.107064

36. Pau, G., Fuchs, F., Sklyar, O., Boutros, M., & Huber, W. (2010). EBImage—An R package for image processing with applications to cellular phenotypes. Bioinformatics, 26(7), 979–981. 10.1093/bioinformatics/btq046

37. Peng, J., Rezaei, E. E., Zhu, W., Wang, D., Li, H., Yang, B., & Sun, Z. (2022). Plant Density Estimation Using UAV Imagery and Deep Learning. Remote Sensing, 14(23). 10.3390/rs14235923

38. Postma, J. A., Hecht, V. L., Hikosaka, K., Nord, E. A., Pons, T. L., & Poorter, H. (2021). Dividing the pie: A quantitative review on plant density responses. Plant, Cell & Environment, 44(4), 1072–1094. 10.1111/pce.13968

39. Pranga, J., Borra-Serrano, I., Quataert, P., De Swaef, T., Vanden Nest, T., Willekens, K., Ruysschaert, G., Janssens, I. A., Roldán-Ruiz, I., & Lootens, P. (2024). Quantification of species composition in grass-clover swards using RGB and multispectral UAV imagery and machine learning. Frontiers in Plant Science, 15. 10.3389/fpls.2024.1414181

40. Rajković, B., Lovrić, A., Maričević, M., Novoselović, D., & Šarčević, H. (2025). Validation of QTLs for Resistance to Pre-Harvest Sprouting in a Panel of European Wheat Cultivars. Plants, 14(9), 1342. 10.3390/plants14091342

41. Rebetzke, G. J., Botwright, T. L., Moore, C. S., Richards, R. A., & Condon, A. G. (2004). Genotypic variation in specific leaf area for genetic improvement of early vigour in wheat. Field Crops Research, 88(2), 179–189. 10.1016/j.fcr.2004.01.007

42. Rebetzke, G. J., & Richards, R. A. (1999). Genetic improvement of early vigour in wheat. Australian Journal of Agricultural Research, 50(3), 291–302. 10.1071/A98125

43. Rodriguez, C., Carlsson, G., Englund, J.-E., Flöhr, A., Pelzer, E., Jeuffroy, M.-H., Makowski, D., & Jensen, E. S. (2020). Grain legume-cereal intercropping enhances the use of soil-derived and biologically fixed nitrogen in temperate agroecosystems. A meta-analysis. European Journal of Agronomy, 118, 126077. 10.1016/j.eja.2020.126077

44. Salomon, J., Enjalbert, J., & Flutre, T. (2026). Joint modeling of social genetic effects in mono-and pluri-specific groups: Case study in intercrops (p. 2026.03.27.714849). bioRxiv. 10.64898/2026.03.27.714849

45. Sampoux, J.-P., Giraud, H., & Litrico, I. (2020). Which Recurrent Selection Scheme To Improve Mixtures of Crop Species? Theoretical Expectations. G3 Genes|Genomes|Genetics, 10(1), 89–107. 10.1534/g3.119.400809

46. Sankaran, S., Khot, L. R., & Carter, A. H. (2015). Field-based crop phenotyping: Multispectral aerial imaging for evaluation of winter wheat emergence and spring stand. Computers and Electronics in Agriculture, 118, 372–379. 10.1016/j.compag.2015.09.001

47. Shen, J., Wang, Q., Zhao, M., Hu, J., Wang, J., Shu, M., Liu, Y., Guo, W., Qiao, H., Niu, Q., & Yue, J. (2024). Mapping Maize Planting Densities Using Unmanned Aerial Vehicles, Multispectral Remote Sensing, and Deep Learning Technology. Drones, 8(4). 10.3390/drones8040140

48. Tilman, D., Cassman, K. G., Matson, P. A., Naylor, R., & Polasky, S. (2002). Agricultural sustainability and intensive production practices. Nature, 418(6898), 671–677. 10.1038/nature01014

49. Varela, S., Dhodda, P. R., Hsu, W. H., Prasad, P. V. V., Assefa, Y., Peralta, N. R., Griffin, T., Sharda, A., Ferguson, A., & Ciampitti, I. A. (2018). Early-Season Stand Count Determination in Corn via Integration of Imagery from Unmanned Aerial Systems (UAS) and Supervised Learning Techniques. Remote Sensing, 10(2). 10.3390/rs10020343

50. Vong, C. N., Conway, L. S., Zhou, J., Kitchen, N. R., & Sudduth, K. A. (2021). Early corn stand count of different cropping systems using UAV-imagery and deep learning. Computers and Electronics in Agriculture, 186, 106214. 10.1016/j.compag.2021.106214

51. Wang, D., Shi, L., Yin, H., Cheng, Y., Liu, S., Wu, S., Yang, G., Dong, Q., Ge, J., & Li, Y. (2025). A Detection Approach for Wheat Spike Recognition and Counting Based on UAV Images and Improved Faster R-CNN. Plants, 14(16), 2475. 10.3390/plants14162475

52. Weiner, J., & Freckleton, R. P. (2010). Constant Final Yield. Annual Review of Ecology, Evolution, and Systematics, 41(Volume 41, 2010), 173-192. 10.1146/annurev-ecolsys-102209-144642

53. Whaley, J. M., Sparkes, D. L., Foulkes, M. J., Spink, J. H., & Et., A. (2000). The physiological response of winter wheat to reductions in plant density. Annals of Applied Biology. 10.1111/j.1744-7348.2000.tb00048.x

54. Wilke, N., Siegmann, B., Postma, J. A., Muller, O., Krieger, V., Pude, R., & Rascher, U. (2021). Assessment of plant density for barley and wheat using UAV multispectral imagery for high-throughput field phenotyping. Computers and Electronics in Agriculture, 189, 106380. 10.1016/j.compag.2021.106380

55. Yang, T., Jay, S., De Solan, B., Li, Y., Zhu, C., Liu, S., & Baret, F. (2025). Estimating small-grain cereal plant density at early growth stages using leaf tip density dynamics derived from submillimeter-scale RGB imagery. Computers and Electronics in Agriculture, 239, 111053. 10.1016/j.compag.2025.111053

56. Yin, H., Yang, H., Hu, Y., Li, F., & Yu, K. (2025). Drone RGB Imagery Color and Texture Information Have Varied Importance in Predicting Potato Aboveground Biomass in Different Growth Stages. Potato Research. 10.1007/s11540-025-09855-z

57. Yu, Y., Stomph, T.-J., Makowski, D., Zhang, L., & van der Werf, W. (2016). A meta-analysis of relative crop yields in cereal/legume mixtures suggests options for management. Field Crops Research, 198, 269–279. 10.1016/j.fcr.2016.08.001

58. Yue, J., Yang, G., Tian, Q., Feng, H., Xu, K., & Zhou, C. (2019). Estimate of winter-wheat above-ground biomass based on UAV ultrahigh-ground-resolution image textures and vegetation indices. ISPRS Journal of Photogrammetry and Remote Sensing, 150, 226–244. 10.1016/j.isprsjprs.2019.02.022

59. Zou, M., Liu, Y., Fu, M., Li, C., Zhou, Z., Meng, H., Xing, E., & Ren, Y. (2024). Combining spectral and texture feature of UAV image with plant height to improve LAI estimation of winter wheat at jointing stage. Frontiers in Plant Science, 14. 10.3389/fpls.2023.1272049

